# Click editing enables programmable genome writing using DNA polymerases and HUH endonucleases

**DOI:** 10.1101/2023.09.12.557440

**Authors:** Joana Ferreira da Silva, Connor J. Tou, Emily M. King, Madeline L. Eller, Linyuan Ma, David Rufino-Ramos, Benjamin P. Kleinstiver

## Abstract

Genome editing technologies that install diverse edits can widely enable genetic studies and new therapeutics. Here we develop click editing, a genome writing platform that couples the advantageous properties of DNA-dependent DNA polymerases with RNA-programmable nickases (e.g. CRISPR-Cas) to permit the installation of a range of edits including substitutions, insertions, and deletions. Click editors (CEs) leverage the “click”-like bioconjugation ability of HUH endonucleases (HUHes) with single stranded DNA substrates to covalently tether “click DNA” (clkDNA) templates encoding user-specifiable edits at targeted genomic loci. Through iterative optimization of the modular components of CEs (DNA polymerase and HUHe orthologs, architectural modifications, etc.) and their clkDNAs (template configurations, repair evading substitutions, etc.), we demonstrate the ability to install precise genome edits with minimal indels and no unwanted byproduct insertions. Since clkDNAs can be ordered as simple DNA oligonucleotides for cents per base, it is possible to screen many different clkDNA parameters rapidly and inexpensively to maximize edit efficiency. Together, click editing is a precise and highly versatile platform for modifying genomes with a simple workflow and broad utility across diverse biological applications.

## Introduction

The discovery and development of CRISPR-Cas enzymes has expanded our ability to make rapidly customizable modifications to the human genome through RNA-programmable technologies^1,2^. Despite these capabilities, the precise modification of DNA sequences via nuclease-based methods that generate DNA double-strand breaks (DSBs) carries risks for undesirable byproducts or consequences, including unwanted insertion or deletion mutations (indels), chromosome-scale changes, p53 activation, and cell-cycle arrest^3–15^. Moreover, applications that rely on homology-directed repair (HDR) impose challenges for cell-cycle dependence^16,17^. Thus, next-generation technologies that install targeted DNA modifications independently of the cell cycle and that don’t require DNA DSBs can overcome these caveats.

Recent efforts have developed DSB-independent technologies capable of nucleotide-level changes. For example, base editors (BEs) are comprised of a DNA nickase (e.g. nCas9) fused to a cytosine or adenine deaminase domain, enabling efficient C-to-T or A-to-G transition substitutions^18–20^. However, BEs are limited in the scope of possible edits they can generate, can be prone to unwanted bystander editing, and have open questions related to on- and off-target DNA and RNA editing^21–25^. Alternatively, polymerase-based DNA writing technologies have been developed for a variety of applications including genetic diversification^26–29^, installing frameshift mutations^30–32^, or writing customized sequence edits^2,33–37^. One class of polymerase-based DNA editors, prime editors (PEs), are comprised of nCas9 fused to a reverse transcriptase (RT) to enable the genetic writing of small edits programmed on a prime editor guide RNA (pegRNA)^34^.

We sought to explore the use of DNA-dependent polymerases (DDPs) for genome editing, given their ubiquitous presence in cells and potentially advantageous attributes. DDPs are capable of high-fidelity polymerization, exhibit high substrate processivity, and are likely to be enzymatically active across nearly any cell type due to high dNTP affinity^38,39^. Furthermore, DDPs are compatible with user-specifiable DNA oligonucleotide (oligo) templates, which may offer advantages in terms of simplicity, scalability, customizability, chemical stability, edit purity, cost, and that oligos are widely used and clinically validated molecules^40^. We therefore envisioned that fusion or recruitment of a DDP to nCas9 (**Fig. 1a**) might create a class of genome writing technologies with valuable distinctions compared to prior approaches. We hypothesized that the use of a single stranded DNA (ssDNA) tethering domain may improve writing efficiency by enabling the localization of the modification-encoding template (**Fig. 1a**). The tripartite ssDNA would include a recognition sequence for a protein or peptide capable of binding nucleic acids, a polymerization template (PT) containing an edit of interest, and a primer binding site (PBS) that bears homology to the target site’s nicked non-target strand (NTS) (**Fig. 1a**).

**Figure 1.**
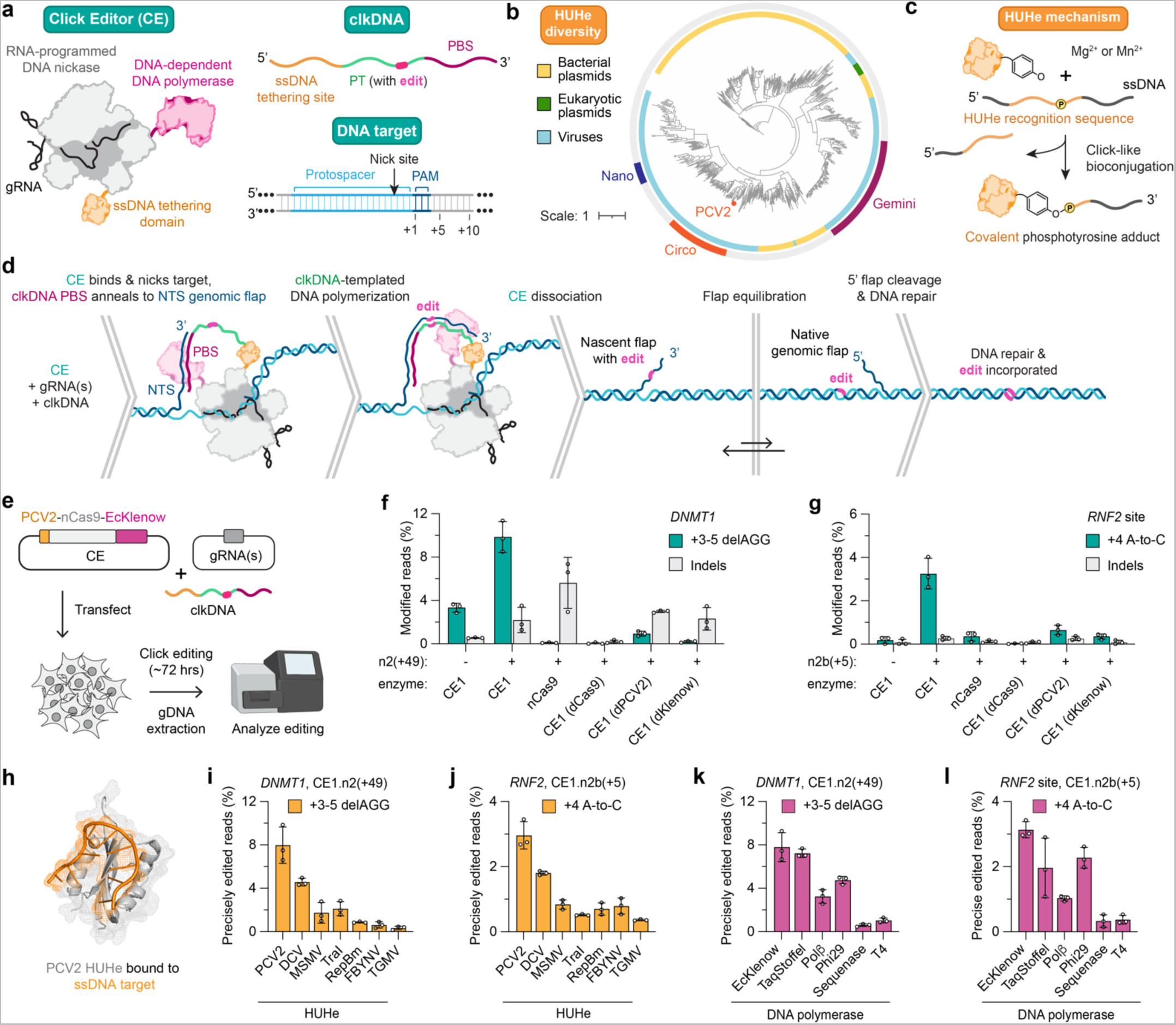
Overview and development of click editing. **a,** Schematic of a click editor (CE), which is a fusion protein consisting of an RNA-programmed DNA nickase, a DNA-dependent DNA polymerase, and a ssDNA tethering domain (e.g. an HUH endonuclease; HUHe) paired with a guide RNA (gRNA). The click-DNA (clkDNA) template is a single-stranded DNA oligonucleotide that encodes a primer binding site (PBS), a polymerase template (PT), and an HUHe recognition site **b,** Phylogenetic tree generated from 709 sequences^47^ depicting a small subset of HUHe diversity across domains of life. Scale represents the fractional distance relatedness between sequences. **c,** Schematic of an HUHe forming a covalent phosphotyrosine adduct with a ssDNA molecule, where the HUHe binds a recognition sequence to initiate a click-like conjugation reaction. **d,** Stepwise click editing mechanism involving: (1) a DNA target site nick to release the non-target strand (NTS) 3’-genomic flap, (2) NTS flap hybridization with the clkDNA PBS, (3) NTS-PBS junction to prime synthesis by the DNA-dependent DNA polymerase, (4) extension of the 3’ NTS flap to polymerize from the edit-encoding PT of the clkDNA, (5) equilibration between the newly synthesized 3’ and native genomic 5’ flaps, and (6) 5’-flap cleavage leading to edit incorporation. **e,** Schematic of click editing transfections in HEK 293T cells, involving co-transfection of a CE plasmid (porcine circovirus 2 (PCV2) HUHe fused to nSpCas9(H840A) and Klenow fragment from *E.coli* DNA polymerase I (D355A, D357A) (EcKlenow)), a clkDNA, and one (or two) gRNA plasmid(s). Editing efficiency is assessed 72 hours post-transfection following genomic DNA extraction and amplicon sequencing. **f,g,** Percentage of sequencing reads with precise edits or reads with insertion or deletion mutations (indels) using the *DNMT1* gRNA and a clkDNA with PBS13-PT12 encoding a +3-5 AGG deletion (with a +49 nick; **panel f**), or the *RNF2* gRNA and a clkDNA with PBS15-PT14 encoding a +4 A-to-C substitution (with a +5 ‘2b’ nick; **panel g**). CE1, CE (PCV2-nSpCas9 (H840A)-EcKlenow) with one gRNA to direct non-target strand nicking; CE1.n2, CE1 with an additional gRNA to direct nicking (i.e. ngRNA) targeted against the non-edited strand at a specified distance from the nick generated by the primary gRNA; CE1.n2b, CE1 with a ngRNA that binds only to the edited strand, directing nicking to the unedited strand; nCas9, CE1.n2 with nCas9 (no HUHe or DNA pol.) and a clkDNA lacking the HUHe recognition site; dCas9, CE1.n2 with a catalytically-deactivated Cas9 (dCas9; D10A, H840A) fused to PCV2 and EcKlenow; dPCV2, CE1.n2 with a catalytically inactive PCV2 (Y96F) fused to an nCas9 and EcKlenow; dKlenow, CE1.n2 with a catalytically inactive EcKlenow (D355A, D357A, D705A, D882A) fused to nCas9 and PCV2. Data in **panels f** and **g** from HEK 293T cell experiments; mean, s.d., and individual datapoints shown for n = 3 independent biological replicates. **h,** Representative structure of the PCV2 HUHe (grey) bound to a ssDNA substrate (orange) (PDB ID: 6WDZ). **i,j,** Percentage of sequencing reads with precise edits when using CE constructs encoding different HUHe domains to install edits using the *DNMT1* or *RNF2* gRNAs (**panels i** and **j**, respectively). DCV, duck circovirus; MSMV, maize striate mosaic virus; TraI, *E.coli* conjugation protein TraI; RepBm, RepB *Fructobacillus tropaeola*; FBYNV, fava bean necrosis yellow virus; TGMV, tomato golden mosaic virus. **k,l,** Percentage of sequencing reads with precise edits when using CE constructs encoding different DNA-dependent DNA polymerases installing edits using the *DNMT1* or *RNF2* gRNAs (**panels k** and **l**, respectively). TaqStoffel, Stoffel fragment from *Thermus aquaticus* DNA polymerase; Polβ, human polymerase beta; Phi29, DNA polymerase from bacteriophage ϕ29 (D169A); Sequenase, engineered truncation of T7 bacteriophage DNA polymerase; T4, T4 bacteriophage DNA polymerase. Data in **panels i-l** from HEK 293T cell experiments; mean, s.d., and individual datapoints shown for n = 3 independent technical replicates.

An ideal ssDNA recruitment domain would have specificity for the provided ssDNA template, be small in coding sequence, have rapid kinetics to catalyze covalent protein-DNA adducts, and not require any specialized and/or expensive modifications. Currently, HUH endonucleases (HUHes) uniquely meet these criteria. HUHes are small proteins spread across all domains of life (**Fig. 1b**) that carry out diverse ssDNA-specific transactions, including ssDNA viral replication, conjugation, transposition, and others^41^. The general class of HUH replication endonucleases and relaxases perform sequence-specific bioconjugation with ssDNA substrates that encode short ~8-40 nt recognition sequences^41^ (**Fig. 1c**). Minimized HUH domains have been used in biological applications as “HUH tags”^42,43^, including as a Cas9 nuclease-based covalent tether for HDR donor templates^44^.

Here we describe the development of click editors (CEs), which leverage the “click-like” protein-substrate attachment biochemistry of HUHes to localize ssDNA oligo template “click DNAs” (clkDNAs) for precision genome editing. CEs can be programmably directed to target sites by gRNAs to install user-specifiable genome modifications encoded on the clkDNA via clkDNA-templated polymerization (**Fig. 1d**). The simplicity of using unmodified DNA molecules as clkDNA templates permits the rapid and inexpensive optimization of editing efficiency by varying parameters including edit type and additional silent mutations, where clkDNAs can be screened in a high-throughput format without requiring molecular cloning steps. We demonstrate that CE enzymes are modular in composition and can utilize various DDP or HUHe domains, identifying opportunities for continued engineering and optimization. Our results highlight potential advantages of DDPs as effectors in the broader class of polymerase-based DNA writing technologies. Together, click editors are capable of a diversity of edit types (all substitutions and short insertions or deletions) for programmable, versatile, and precise HDR- and DSB-independent DNA modification with broad potential for diverse biological applications.

## Results

We constructed an initial CE fusion protein (CE1) that combined the HUHe from porcine circovirus 2 (PCV2) and a 3’-5’ exonuclease-deficient Klenow fragment from *E. coli* DNA polymerase I (EcKlenow^45^) with an SpCas9 nickase (nCas9; H840A) (**Figs. 1a,d,e**). CEs initiate NTS nicking to release the endogenous genomic flap^46^ while covalently tethering a clkDNA template (encoding a PT harboring the desired edit and a PBS) to the target site via the HUHe domain (**Fig. 1d**). Annealing of the tethered clkDNA PBS to the nicked NTS creates a primed substrate for clkDNA-templated DNA polymerization by the CE-fused DDP, resulting in an extended 3’ flap containing the desired edit. Subsequent flap equilibration and DNA repair to incorporate the nascent 3’ flap leads to precise installation of the edit at the target site (**Fig. 1d**). Transfections were performed using separate CE and single gRNA expression plasmids along with a clkDNA (encoding the 13 nt PCV2 recognition sequence^44^), an approach that we termed CE1 or CE1.n1 due to use of a single primary gRNA (where CE1 defines the CE enzyme and n1 defines the gRNA/nicking strategy). The clkDNAs encoded a 3-bp deletion at *DNMT1* or an A-to-C transversion at *RNF2* (**Figs. 1f** and **1g**, respectively) and were modified with two 3’ phosphorothioate (PS) linkages. Amplicon sequencing from experiments using CE1 revealed 3.33% and 0.18% precise editing at *DNMT1* and *RNF2*, respectively, with minimal indel byproducts (**Figs. 1f-g**).

To improve edit efficiency and explore CE dependencies, we tested various gRNA and CE configurations. Similar to BEs^18^ and PEs^34^ a secondary nick on the opposite strand should bias DNA repair to preferentially utilize the edited strand as the correct template. We performed experiments with CE1 and a secondary gRNA to direct CE-mediated nicking (ngRNA), leading to CE1.n2 or CE1.n2b conditions where the ngRNA is either distal from or overlapping the intended edit, respectively (**Sup. Fig. 1**). With CE1.n2 we observed ~3-fold increase in precise editing compared to CE1 at *DNMT1*, achieving 9.85% editing (**Fig. 1f** and **Sup. Fig. 3a**); a clkDNA titration using these conditions revealed that 16 pmol of clkDNA led to optimal editing efficiency in our initial experiments (**Sup. Fig. 2a, Sup. Note 1**). When using an n2b approach at *RNF2* (where the ngRNA overlaps the intended edit), we observed a ~18-fold increase in editing versus CE1, reaching 3.26% (**Fig. 1g** and **Sup. Fig. 3b**). Importantly, transfecting various control conditions nearly or fully abolished click editing, when using plasmids encoding nCas9 only (no DDP or HUHe fusions) or CE1.n2 containing catalytically inactive Cas9 (no NTS nick with HUH-less clkDNA), inactive PCV2 (diminished clkDNA recruitment), or inactive Klenow (attenuated polymerization) (**Figs. 1f,g**, and **Sup. Note 2**). Notably, the n2b-mediated indels for *RNF2* were lower compared to n2-induced indels at *DNMT1*, consistent with the hypothesis that ngRNAs that bind only after editing has occurred reduces the co-occurrence of nicks and the subsequent generation of DSBs (analogous to the PE3b strategy^34^) (**Figs. 1f,g**, and **Sup. Note 2**). These experiments demonstrate that CEs can achieve precise genome edits and that all components (nCas9, DDP, HUHe, and clkDNA) are required for effective click editing.

Next, we explored alternative HUHe and DDP enzymes in our modular CE architecture. Given the diversity of HUHes (**Fig. 1b**), we tested a variety of domains involved in the replication of circoviruses, geminiviruses and nanoviruses, and the conjugative relaxase TraI^42^. At both *DNMT1* and *RNF2*, CE1.n2 editing was the most efficient with our original CE1 construct containing the PCV2 HUHe fused to nCas9 (**Figs. 1h-j** and **Sup. Figs. 4a,b**). We then explored whether the use of different family A, B, or X DDP enzymes might alter click editing efficiency. Again, we found that our original design containing EcKlenow was the most efficient at the *DNMT1* and *RNF2* sites, although the Stoffel fragment of *Thermus aquaticus* DNA polymerase (TaqStoffel^48^) and bacteriophage Phi29 DNA Polymerase (Phi29^49^; D169A for exonuclease inactivation^50^) also exhibited nearly comparable precise editing (**Figs. 1k,l** and **Sup. Figs. 4c,d**). Furthermore, decreasing the clkDNA dosage from 16 to 12 pmol increased editing efficiency (**Sup. Fig. 2b** and **Sup. Note 1**), potentially by decreasing gRNA sequestration (from excess clkDNA interactions with the gRNA spacer^51,52^) and/or by reducing potential cleavage of the gRNA by RNAseH due to the RNA:DNA duplex. Additional experiments testing alternative linkers between PCV2 and nCas9, or truncations of a flexible C-terminal region of PCV2, did not improve editing efficiency (**Sup. Fig. 4e**). Therefore, we proceeded with the PCV2-nCas9-EcKlenow CE construct for further characterization.

We hypothesized that click editing efficiency might be improved by testing additional parameters of clkDNA design. For instance, optimal annealing of the PBS to the genomic NTS flap could be crucial to form a stable template-primer junction for polymerase initiation, or the length of the PT could influence flap-genome hybridization and/or flap equilibration (**Fig. 1d**). The use of HUHes uniquely permits the rapid and high-throughput assessment of clkDNA properties, since HUHes form covalent protein-clkDNA adducts with simple unmodified ssDNA oligos (without requiring chemical or specialized modifications; **Fig. 1c**). To implement clkDNA optimizations, pre-normalized 96-well plates of simple unmodified DNA oligos can be purchased at relatively low cost, a rapid process that does not require additional cloning steps (**Fig. 2a**).

**Figure 2.**
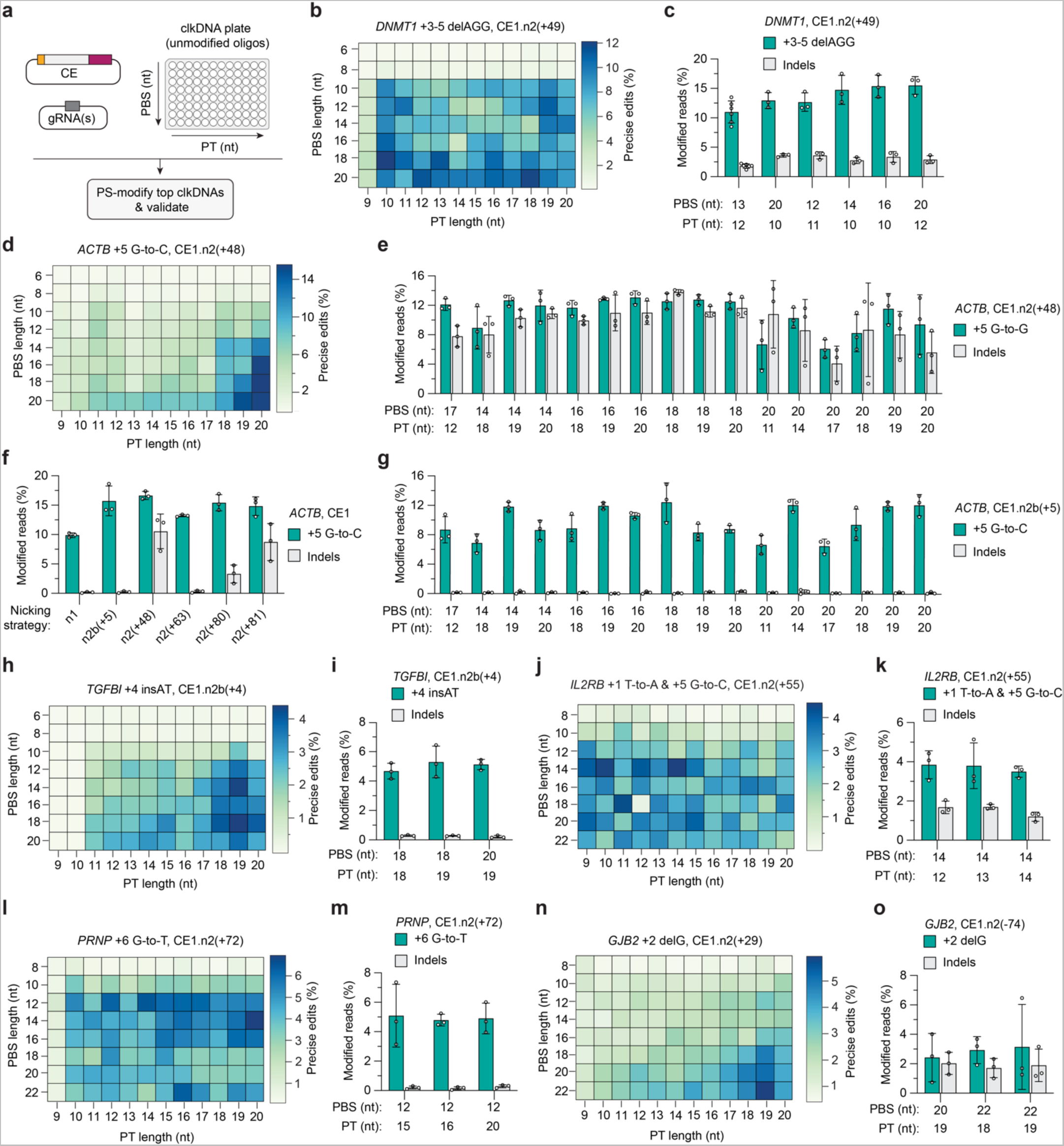
Optimization of clkDNA parameters. **a,** Schematic of clkDNA screens in 96-well format. The CE, gRNA and ngRNA (CE1.n2) are transfected together with up to 96 unprotected clkDNA oligonucleotides (oligos) with various PBS and PT lengths arrayed on a plate. Optimal clkDNA candidates can then be further chemically modified (e.g. with two phosphorothioate (PS) linkages) for validation studies. **b,** Percentage of sequencing reads with a precise +3-5 AGG deletion using the *DNMT1* gRNA, in a clkDNA screen using unmodified oligos to vary the PBS and PT lengths. **c,** Percentage of sequencing reads with precise edits or indels, when assessing the most efficient *DNMT1* clkDNAs but with 2×3’-PS linkages on the clkDNA. **d,** Percentage of sequencing reads with a precise +5 G-to-C transversion using the *ACTB* gRNA, in a clkDNA screen using unmodified oligos to vary the PBS and PT lengths. **e,** Percentage of sequencing reads with precise edits or indels, when assessing the most efficient *ACTB* clkDNAs but with 2×3’-PS linkages on the clkDNA. **f,** Percentage of sequencing reads with precise edits or indels with different nicking gRNAs (ngRNA) targeting *ACTB* and a 2×3’-PS protected clkDNA of PBS16-PT19. ***g***, Percentage of sequencing reads with precise edits or indels, when assessing the most efficient *ACTB* clkDNAs with 2×3’-PS linkages on the clkDNA and a 2b ngRNA (n2b, +5). **h,** Percentage of sequencing reads with a precise +4 AT insertion using the *TGFBI* gRNA, in a clkDNA screen using unmodified oligos to vary the PBS and PT lengths. **i,** Percentage of sequencing reads with precise edits or indels, when assessing the most efficient *TGFBI* clkDNAs but with 2×3’-PS linkages on the clkDNA. **j,** Percentage of sequencing reads with a precise dual +1 T-to-A & +5 G-to-C edit using the *IL2RB* gRNA, in a clkDNA screen using unmodified oligos to vary the PBS and PT lengths. **k,** Percentage of sequencing reads with precise edits or indels, when assessing the most efficient *IL2RB* clkDNAs but with 2×3’-PS linkages on the clkDNA. **l,** Percentage of sequencing reads with a precise +6 G-to-T edit using the *PRNP* gRNA, in a clkDNA screen using unmodified oligos to vary the PBS and PT lengths. **m,** Percentage of sequencing reads with precise edits or indels, when assessing the most efficient *PRNP* clkDNAs but with 2×3’-PS linkages on the clkDNA. **n,** Percentage of sequencing reads with a precise +2 G deletion using the *GJB2* gRNA, in a clkDNA screen using unmodified oligos to vary the PBS and PT lengths. **o,** Percentage of sequencing reads with precise edits or indels, when assessing the most efficient *GJB2* clkDNAs but with 2×3’-PS linkages on the clkDNA. Data in **panels b**,**d**,**h**,**j**,**l**,**n** from HEK 293T cell experiments; mean, s.d., and individual datapoints shown for n = 3 independent biological replicates. Data in **panels c**,**e**,**f**,**g**,**i**,**k**,**m**,**o** from HEK 293T cell experiments; mean, s.d., and individual datapoints shown for n = 3 independent technical replicates.

To scalably assess clkDNA parameters, we ordered and screened 96 clkDNA configurations including combinations of PBSs from 6-20 nucleotides (nt) and PTs from 9-20 nt. We initially tested this approach for the *DNMT1* +3-5 AGG deletion, which previously yielded nearly 10% precise editing when using a 13 nt PBS and a 12 nt PT clkDNA (PBS13-PT12) (**Fig. 1f**). Experiments to test all 96 unmodified clkDNA oligos yielded editing with several clkDNAs up to ~12% (**Fig. 2b** and **Sup. Figs. 5a-c**). We then performed a validation experiment by selecting 15 clkDNAs that yielded higher efficiencies in the primary screen (**Sup. Fig. 5c**) and testing them with two chemically modified 3’ phosphorothioate (PS) linkages which should improve clkDNA stability (**Sup. Figs. 5d,e**). We observed good correlation between the efficiencies observed with unmodified and PS-modified clkDNAs (**Sup. Figs. 5d,e**), reaching up to 15.4% precise editing with a PBS16-PT10 modified clkDNA (**Fig. 2c**) and leading to >50% improvement in efficiency compared to the initial unmodified clkDNA (**Fig. 1f**).

We then explored the generalizability of our scalable clkDNA optimization across other new sites to install various edits. Using an *ACTB*-targeted gRNA and 96 different clkDNAs encoding a G-to-C transversion, we observed a distinct trend towards higher efficiencies with longer PT and PBS lengths, reaching up to 15.5% precise editing with the PBS20-PT20 clkDNA (**Fig. 2d**). However, we also observed high levels of indels when using the n2(+48) ngRNA (**Fig. 2e, Sup. Figs. 6a-c**), motivating us to explore additional ngRNAs. Among the CE1 condition with no ngRNA, the +48 ngRNA, and four additional ngRNAs, the n2b(+5) strategy in combination with an optimal clkDNA configuration (PBS16-PT19) yielded up to 15.74% mean precise editing with minimal indels (**Fig. 2f-g**). These results highlight how careful ngRNA and clkDNA selection can dramatically improve edit efficiency and purity, parameters that are easily optimized with click editing.

Following a similar approach, we then performed clkDNA screens at additional genomic sites. We achieved nearly 5% precise editing at *TGFBI* for a +4 AT insertion (**Figs. 2h,i**, and **Sup. Figs. 7a-d**), at *IL2RB* for dual substitutions (**Figs. 2j,k**, and **Sup. Figs. 8a-d**), at *PRNP* for a +6 G-to-T transversion (**Figs. 2l,m**, and **Sup. Figs. 9a-d**), and at *GJB2* for a +2 G deletion (**Fig. 2n,o** and **Sup. Figs. 10a-d**). Using over 600 oligos across these six distinct clkDNA screens to install various edits at multiple genomic sites and using several ngRNA types, we observed some common clkDNA parameter trends, including that PBSs <10 nt do not generally support productive click editing with our current CE configuration, and that in most cases longer PBSs and PTs are typically more effective (**Figs. 2b-2o** and **Sup. Figs. 5-10**). Among these sites and edits, precise click editing using parameter-optimized clkDNAs ranged from 3.50% to 15.74% with minimal indels. Together, the results from our clkDNA optimizations demonstrate that highly precise editing can be achieved through facile and scalable clkDNA and ngRNA screening.

Prior studies suggested that inhibition or evasion of DNA repair (e.g. mismatch repair; MMR) can improve prime editing efficiencies by counteracting excision of the edited flap^53,54^. Design of strategies that install additional edits (e.g. silent substitutions) along with the desired edit can suppress recognition of the installed mismatch, thereby increasing editing efficiencies (**Fig. 3a**). Since the resolution of nascent 3’ DNA flaps encoding mismatches installed by click or prime editing likely proceed through similar repair mechanisms, we investigated whether varying the base composition of the clkDNA PT could enhance repair evasion to improve precise click editing. We selected a clkDNA to install a dual T-to-A and G-to-C edit at the *IL2RB* locus, for which we previously achieved nearly 4% precise editing with the PBS14-PT14 clkDNA and CE1.n2 (**Fig. 2j**). In editing experiments with CE1 and CE1.n2 and various clkDNAs encoding additional silent substitutions, we observed 24.7- and 4.1-fold increases in precise click editing, respectively, with the most effective clkDNA design containing 2 additional silent substitutions compared to the original design (**Fig. 3b**). We then designed six clkDNAs for a new edit to install a G-to-T transversion at the *VEGFA* locus, with various combinations of five additional substitutions (**Fig. 3c**). Compared to 12.4% precise editing when using CE1 and a clkDNA encoding only the primary edit, we observed an average of 26.6% editing (with 2.7% indels) using a clkDNA encoding three additional repair-evading substitutions (a 2.2-fold increase; **Fig. 3c** and **Sup. Figs. 11a-11d**).

**Figure 3.**
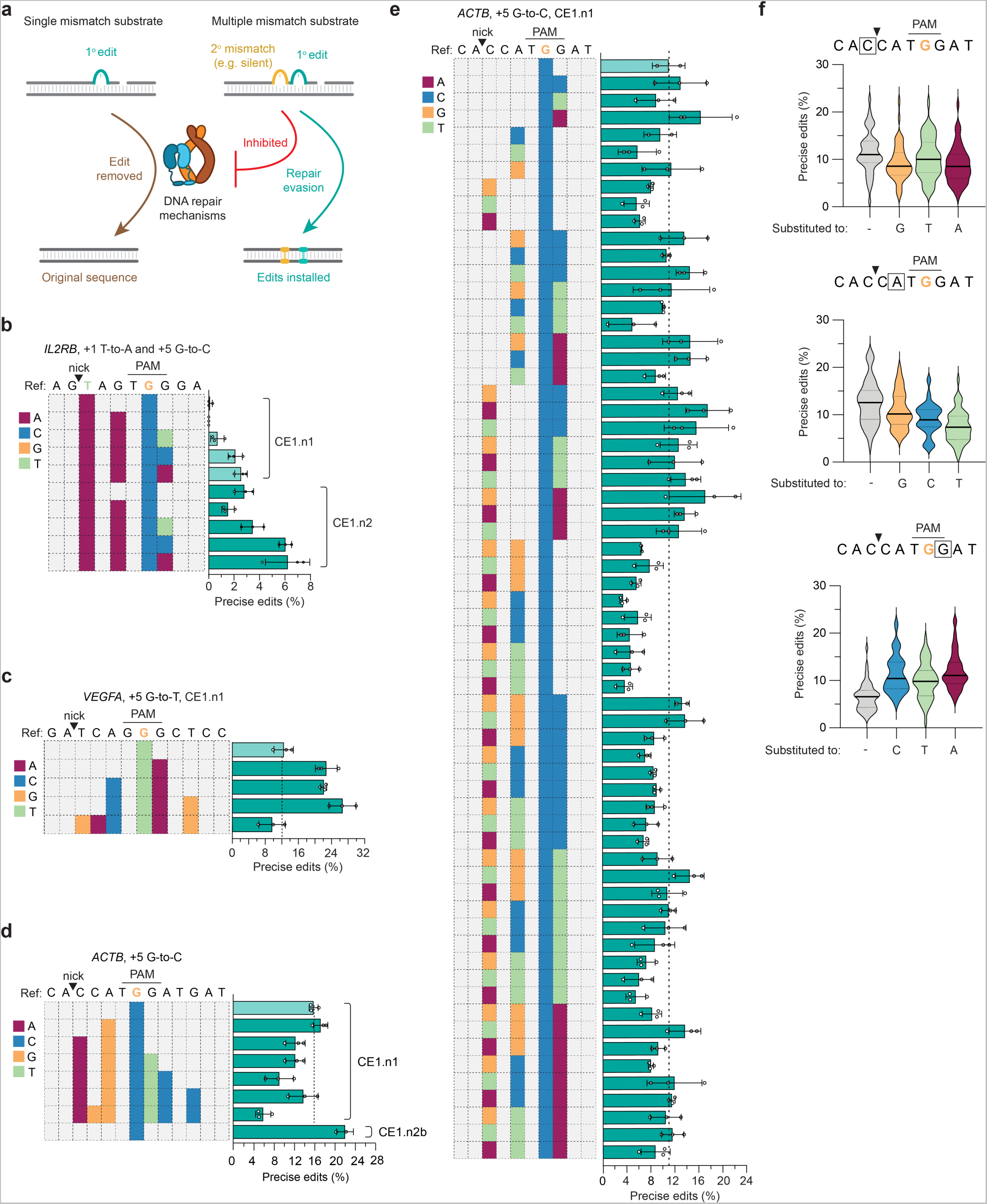
DNA repair evasion through clkDNA modification. **a,** Schematic of DNA repair engagement on substrates with different compositions of mismatches. DNA repair mechanisms can excise the DNA flap encoding the intended edit (1° edit, teal); encoding additional substitutions (2° mismatch, yellow) adjacent to the intended edit (1° edit, teal) may evade excision of the intended edit. **b,** Percentage of sequencing reads with precise +1 T-to-A and +5 G-to-C transversions using CE1.n1 or CE1.n2, the *IL2RB* gRNA, and clkDNAs encoding additional mutations for repair evasion. Colors represent nucleotide changes. Ref:, reference amplicon; triangle, gRNA nick site; PAM, protospacer adjacent motif. **c,** Percentage of sequencing reads with a precise +5 G-to-T transversion using CE1.n1, the *VEGFA* gRNA, and clkDNAs encoding additional mutations for repair evasion. **d,** Percentage of sequencing reads with a precise +5 G-to-C transversion using CE1.n1, the *ACTB* gRNA, and clkDNAs encoding additional mutations for repair evasion. The CE1.n2b condition also has a +5 ngRNA that overlaps the installed mutation. **e,** Percentage of sequencing reads with a precise +5 G-to-C transversion using CE1.n1, the *ACTB* gRNA, and clkDNAs encoding all possible bases in three positions of the clkDNA for repair evasion. **f,** Violin plots depicting percentage of reads with precise edits in *ACTB* depending on the nature and the position of the mutation within the clkDNA. The query base is shown with a box. Data in **panels c**,**d**,**e**,**f** from HEK 293T cell experiments; mean, s.d., and individual datapoints shown for n = 3 independent biological replicates. Data in **panel b** from HEK 293T cell experiments; mean, s.d., and individual datapoints shown for n = 3 independent technical replicates.

Given the improvement in click editing efficiencies that we observed with clkDNAs harboring additional substitutions for the *IL2RB* and *VEGFA* edits, we then took a similar approach for the *ACTB* edit. However, our initial *ACTB* clkDNA designs bearing additional substitutions did not substantially increase precise editing (**Fig. 3d**). Considering that the identity of the additional substitutions(s) may bias mismatch excision (**Fig. 3a**), we sought to leverage the scalability of clkDNA synthesis and screening to test a larger and more diverse set of mismatch-harboring clkDNAs. We tested a total of 64 clkDNAs containing all possible combinations of bases in three specific positions within the *ACTB* clkDNA, along with the +5 G-to-C edit. Our screen yielded clkDNAs with diverse impacts on editing efficiency (**Fig. 3e**) and provided insight into which positions and types of modifications within the clkDNA are favorable or detrimental to precise editing (**Fig. 3f**). This style of experiment again highlights the scalability of CEs, while also demonstrating how click editing may be utilized to study biological processes like repair of DNA mismatches. Future studies to extend this approach across other loci may provide more generalizable knowledge into the types and positions of mismatches that maximize edit efficiency.

Next, we sought to compare click and prime editing. Since CE1 utilizes an unevolved wild-type (WT) EcKlenow polymerase, we included both PE1 and PE2 enzymes in our comparison (which contain a WT or an engineered MMLV reverse transcriptase domain, respectively^34^). The PE2 enzyme combined with an additional ngRNA to direct PE2 nicking is referred to as PE3^34^. We performed experiments using CEs with gRNAs and clkDNAs and PEs with previously optimized pegRNAs^34^, targeting *VEGFA* with no ngRNA, *DNMT1* with an n2(+49) ngRNA, and *ACTB* with an n2b(+5) ngRNA. We observed higher or similar precise editing efficiencies with CE1 compared to PE1, suggesting a similarity in systems when both strategies utilize WT polymerases (**Figs. 4a-c** and **Sup. Fig. 12a**). Compared to PE3 at the *DNMT1* or *ACTB* target sites, or to PE2 at *VEGFA*, editing with CE1 was generally less efficient (though was higher in one instance; **Sup. Fig. 12a**). These results are likely attributable to the higher efficiency endowed by the engineered reverse transcriptase domain in PE2^34^, suggesting that future efforts to engineer EcKlenow or other DDPs could substantially increase CE efficiency.

**Figure 4.**
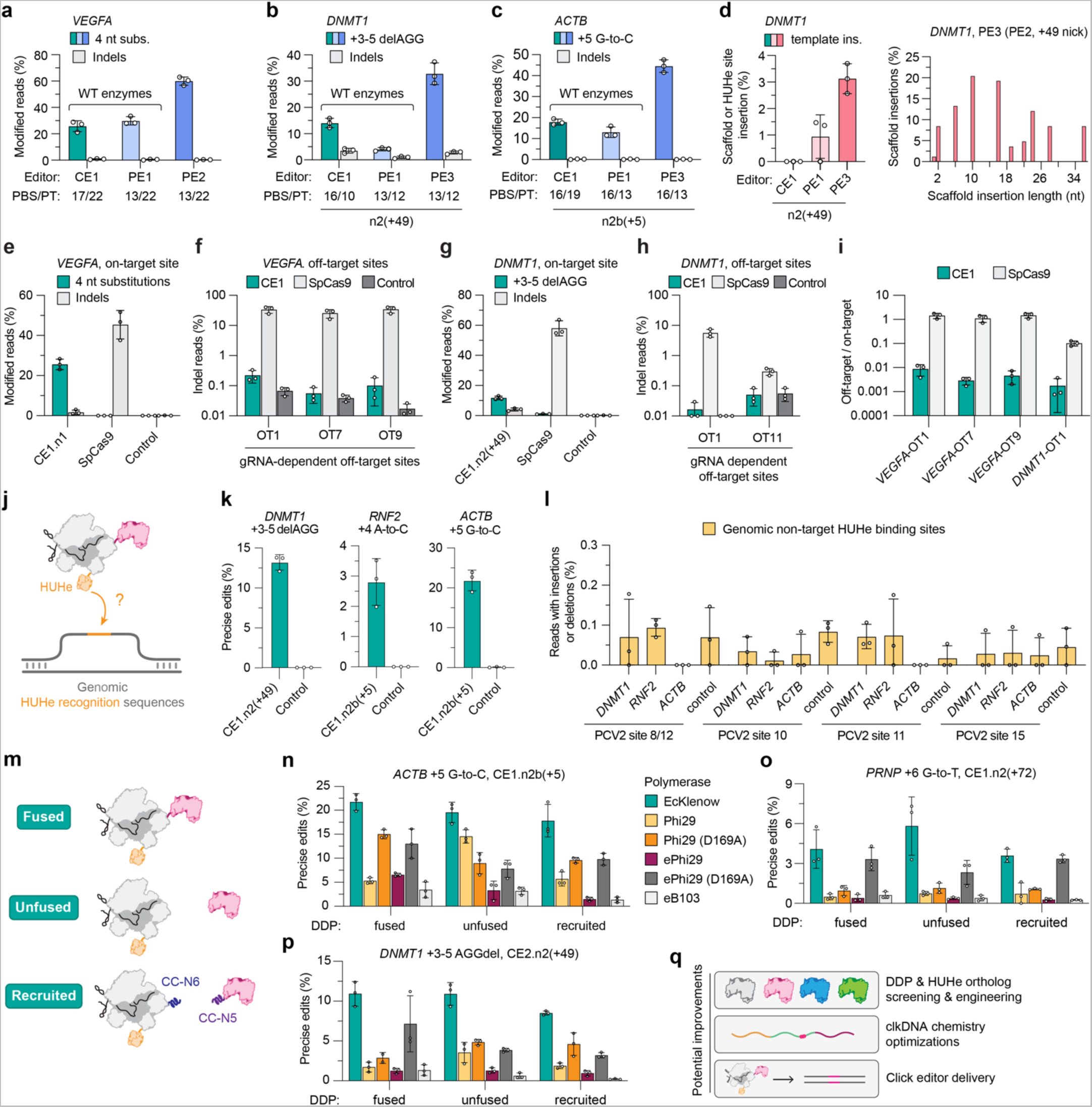
Comparison to prime editing, off-target analyses, and architectural alterations. **a-c,** Percentage of sequencing reads with precise edits or indels using CE1 (PCV2-nSpCas9(H840a)-EcKlenow), PE1 (nSpCas9(H840A)-M-MLV-RT), PE2 (nSpCas9(H840A)-M-MLV-RT(D200N/L603W/T330P/T306K/W313F)^34^), or PE3 (PE2 + ngRNA) when targeting *VEGFA* (with CE1.n1, no ngRNA for CEs or PEs; **panel a**), *DNMT1* (with CE1.n2(+49), using the +49 ngRNA for CEs and PEs; **panel b**), or *ACTB* (with CE1.n2b(+5), using the +5 ngRNA for CEs and PEs; **panel c**). For CEs, clkDNAs were optimized in this study; for PEs, pegRNAs were previously optimized for *VEGFA* and *DNMT1*^34^, and we performed a small optimization of pegRNAs for *ACTB* (see **Sup. Fig. 12b**). WT, wild-type. **d,** Percentage of edited reads at the *DNMT1* locus that contain insertion of the PCV2 HUHe recognition sequence (for CEs; no reads) or the pegRNA scaffold sequence (for PEs) (**left panel**), and distribution of pegRNA scaffold insertion lengths from the PE3 condition (PE2 with +49 nicking gRNA; **right panel**). Reads containing insertions quantified as described in the Methods section. **e,f,** Percentage of reads in experiments using the *VEGFA* gRNA with precise editing or indels at the on-target site (**panel e**) or off-target sites (**panel f** and **Sup. Fig. 13c**) using CE1.n1 or SpCas9 nuclease compared to an untransfected control. **g,h,** Percentage of reads in experiments using the *DNMT1* gRNA with precise editing or indels at the on-target site (**panel g**) or off-target sites (**panel h** and **Sup. Fig. 13d**) using CE1.n2(+49) or SpCas9 nuclease compared to an untransfected control. **i,** Ratio of off-target to on-target editing for selected off-target sites in *VEGFA* and *DNMT1*, using CE1.n1 or CE1.n2, respectively, or SpCas9 (data from **panels f** and **h**). **j,** Schematic of possible HUHe-dependent interaction with genomic sites containing an HUHe recognition sequence that are transiently ssDNA during cellular replication or transcription. **k,** Percentage of sequencing reads with precise edits for *DNMT1, RNF2* and *ACTB* (on-target editing) from experiments with various CE1 conditions. **l,** Percentage of sequencing reads with insertions or deletions at PCV2 HUHe pseudosites in the human genome in various CE1 conditions targeting either *DNMT1*, *RNF2* or *ACTB*. **m,** Schematic of different CE1 architectures tested. CC, coiled-coil domains N5/N6^64,65^; EcKlenow, Klenow fragment from *E.coli* DNA polymerase I (D355A, D357A); Phi29, DNA polymerase from bacteriophage <29; Phi29 (D169A), 3’-5’ exonuclease-deficient Phi29 DNA polymerase; ePhi29, engineered thermostable Phi29 DNA polymerase (M8R, V51A, M97T, G197D, E221K, Q497P, K512E, F526L); ePhi29 (D169), 3’-5’ exonuclease-deficient ePhi29 (D169A, M8R, V51A, M97T, G197D, E221K, Q497P, K512E, F526L); eB103, engineered thermostable B103 DNA polymerase (a Phi29 ortholog) (H73R, A147K, R221Y, A318G, M339L, E359D, K372E, F383L, D384N, A503M, I511V, R544K, T550K). **n-p,** Percentage of sequencing reads with edits in experiments targeting *ACTB*, *PRNP* and *DNMT1* (**panels n-p**, respectively) when using CE1.n2 constructs encoding different DNA-dependent polymerases and different construct architectures. **q,** Potential future optimizations for engineering improved CEs. Data in **panels a**-**i**,**k**,**l** from HEK 293T cell experiments; mean, s.d., and individual datapoints shown for n = 3 independent biological replicates. Data in panels **n-p** from HEK 293T cell experiments; mean, s.d., and individual datapoints shown for n = 3 independent technical replicates.

When comparing CEs and PEs, we also analyzed unwanted insertion mutations at the on-target site. Since the pegRNA is a fusion of the PBS/RTT (reverse transcriptase template) with the gRNA scaffold, the RT domain of PEs can install unwanted gRNA scaffold bases into the target site^34,53,55–58^ (**Sup. Fig. 12b**). With PE-treated samples, we observed >3% pegRNA scaffold incorporation at the *DNMT1* target site (**Fig. 4d** and **Sup. Fig. 12c**). In contrast, no detectable gRNA scaffold or HUHe site incorporation was detected in click edited samples (**Fig. 4d**), suggesting two potential mechanisms to limit unwanted insertions: (1) the inherent separation of template from gRNA in CEs, and (2) covalent attachment of the HUHe to the clkDNA, where the minimal remaining HUHe target site post catalysis on the clkDNA is incompatible with extension due to the HUHe protein footprint^43^ (**Sup. Fig. 12b**). Thus, the CE architecture may be more likely to generate precise edits without unwanted template insertions compared to prime editing with pegRNAs.

Like the high fidelity mechanism of prime editing^59–62^, click editing also requires several proof-reading steps that may reduce the likelihood of gRNA-dependent off-target editing, including pairing of the gRNA spacer with the genomic target site, clkDNA PBS annealing to the NTS, annealing of the nascent 3’ flap to the genomic locus for edit resolution, and use of a nickase instead of a nuclease. To investigate potential off-target edits when using CEs, we targeted a CE or Cas9 nuclease to three target sites in the *VEGFA, DNMT1*, and *ACTB* loci (**Figs. 4e,g**, and **Sup. Fig. 13a**, respectively). We simultaneously assessed on-target editing and potential off-target editing at putative off-target sites closely related in sequence to the on-target sites^63^ (see *Methods;* **Figs. 4e-i** and **Sup. Figs. 13a-13d**). Across 29 off-target sites, when using SpCas9 nuclease we observed considerable indels at three *VEGFA* off-target sites (33.9%, 25.6%, 34.2%) and two *DNMT1* off-target sites (5.7%, 0.3%) (**Figs. 4f,h**, and **Sup. Figs. 13b-13d**). With CEs, we observed dramatically lower near-background levels of off-target indels and no evidence of precise off-target edit installation, supporting a potential high-fidelity mechanism of click editing (**Figs. 4f,h**, and **Sup. Figs. 13b-13d**).

Since HUHes are DNA endonucleases, we also investigated whether spurious interaction of PCV2 HUHe with genomic ssDNA could result in undesired indels at endogenous PCV2 pseudosites (**Fig. 4j**). We identified 16 PCV2 consensus sites in the human genome near NGG protospacer-adjacent motifs (PAMs) (**Sup. Note 3**). Using CE-treated samples containing *DNMT1-*, *RNF2-*, or *ACTB*-targeted gRNAs and clkDNAs (resulting in efficient click editing at the intended on-target sites; **Fig. 4k**), we amplified and sequenced four PCV2 genomic sites, which revealed no elevation in indels compared to untreated control samples (**Fig. 4l**). When we intentionally induced stable R-loops and accessible ssDNA at the PCV2 target sites in a reporter assay (using the CE construct and gRNAs targeted to the PCV2 sites; **Sup Fig. 14a**), we observed slightly elevated HUH-mediated indels at these four sites (**Sup. Figs. 14b-e** and **Sup. Note 3**). Together, these results demonstrate that CE-fused HUHe enzymes are specific towards the HUHe ssDNA recognition sequence encoded on the clkDNA and carry little risk of genomic off-targets, but also suggest that rare occurrences of gRNAs with spacers matching the HUHe sequence should be avoided.

Lastly, we performed a CE architecture optimization to determine the importance of DDP fusion to nCas9, and that also included alternative DDPs from Phi29 and Phi29-like phages given their inherently high fidelity and processivity. While we previously observed slightly lower click editing efficiencies in our initial CE constructs when testing Phi29 DDP compared to EcKlenow CEs (**Figs. 1k,l**, and **Sup. Figs. 4c,d**), we recognized that our previous N-terminal fusion could be detrimental to polymerization activity^66–68^, and that Phi29 is optimally active at 30 °C rather than 37 °C^66,67^. To explore whether Phi29 or similar polymerases could support click editing, we tested wild-type Phi29, an engineered thermostable Phi29 (ePhi29)^69^, and an engineered thermostable Phi29 ortholog (eB103)^70^ in fused, unfused, or polymerase recruited CE architectures (the latter via N5/N6 coiled-coil domains^64,65^) (**Fig. 4m**). Analysis of precise editing at *DNMT1*, *ACTB*, *PRNP, IL2RB and RNF2* revealed optimal activities with our original EcKlenow polymerase, which demonstrated comparable efficiencies across each of the three nCas9-DDP configurations (**Figs. 4n-p** and **Sup. Figs. 15a,b**). Phi29 also permits click editing, with generally increased efficiencies in the unfused configuration and with exonuclease inactivation (D169A). The previously engineered thermostable ePhi29 or B103 mutants did not lead to increased editing efficiencies. Indels varied across configurations, with generally higher rates in the unfused configurations and when using ePhi29 (**Sup. Figs. 15c-15g**). Given that most of these comparisons were performed using clkDNAs resulting from our screens using EcKlenow (aside from the *RNF2* clkDNA which did not undergo a screen), it is possible that alternative polymerases may differ in optimal clkDNA parameters. Together, these results demonstrate that the modularity of the CE complex can enable productive click editing with various polymerases and architectures.

## Discussion

Here we developed click editing, which enables ‘click-to-install’ genome writing via HUHe-mediated covalent clkDNA localization to a target site for DNA-dependent polymerization. CEs can install a diverse range of precise and pure genome edits, with simple design parameters that result in little barrier to entry for new users. The use of simple clkDNA oligo templates facilitates the rapid and scalable interrogation of optimal parameters to maximize efficiency (e.g. PT/PBS lengths and repair evading substitutions), without the need for cost-prohibitive or specialized chemical modifications, RNA or hybrid nucleic acid templates, or additional cloning steps. Indeed, in our study we screened nearly 1,000 clkDNA templates, a feat not financially feasible for other genome editing platforms like prime editing (due to pegRNA synthesis or cloning costs) or nucleases with HDR (due to the expense of long template production). The use of unmodified oligos permits facile initial clkDNA determinant screening, which can then be used as starting point for further chemical modification to improve clkDNA stability, minimize potential immune responses, reduce PBS/spacer interactions between the clkDNA and gRNA, or mitigate potential RNaseH degradation. A more thorough investigation into how optimal clkDNA parameters are preserved or altered across different cells or organisms will provide insight into the generalizability of CEs, an approach that should be facilitated by simplicity of clkDNA synthesis.

There is a vast diversity of HUHe and DDPs that can be explored and leveraged to engineer a suite of CE constructs with improved properties. For instance, other HUHe and DDP orthologs may harbor desirable characteristics including smaller coding sequences, higher efficiency, higher processivity, higher fidelity, faster kinetics, etc. Our initial experiments with CEs compared favorably to the efficiencies observed with the original unengineered MMLV RT domain used in the first generation PE1 construct. These results suggest that the DDP component of current CEs would be an ideal target for improvement, motivating further efforts to engineer DDPs as has been done with PEs (achieving large boosts in efficiency via RT engineering of PE1 to later generation PEs^34,53,55^). Given the modularity of CEs, potential synergistic optimization of each component (DDP, HUHe, nCas9, and clkDNA) suggests a high ceiling in terms of efficiency, particularly since current CEs utilize wild-type EcKlenow and standard oligos (or those with default PS linkage configurations). We envision that these future improvements should offer large benefits for sites where we observed low click editing efficiencies, including the use of alternate polymerases that have different initiation sequence preferences^71^. The CE architecture could also be well-suited to use with other miniature RNA-guided nickases (e.g. Cas9 orthologs^72^ or IscB^73^) where a smaller coding sequence would facilitate improved component manufacturing and delivery due to reduced size.

The use of DDPs in CEs may offer advantages over RT domains in PEs, since DDPs are typically higher fidelity, are capable of longer polymerization processivity, are less sensitive to dNTP availability (which can be low in different cell types and phases of the cell cycle), and given their ubiquitous expression in cells they may be less likely to cause unwanted genome-scale changes (e.g. potential non-specific cDNA production from endogenous RNA transcripts with RTs)^38,39,74^. Furthermore, our data and that of others^34,53,55–58^ demonstrates that reverse transcription into the pegRNA scaffold by the PE RT domain can lead to problematic incorporation of unwanted sequences into the genome, resulting in lower edit purity. We did not observe any clkDNA template/scaffold insertions into the genome, likely because the HUHe protects the remaining few bases of its own binding site post-cleavage, leaving only the writing template of the clkDNA accessible for polymerization. Additionally, efforts to investigate various pegRNA parameters affecting efficiency and purity are laborious and costly, often making the optimization of PEs difficult for routine use. With CEs, we demonstrate the simplicity of screening many different clkDNA configurations and how repair evading mutations can also be easily screened and implemented to enhance edit outcomes (in some cases >24-fold). Finally, the production of pure and high-yield synthetic clkDNA templates offers potential advantages compared to synthetic pegRNAs in non-viral delivery applications, where chemical synthesis of long RNA molecules can be limiting from a purity and cost perspective.

Together, our results highlight the advantageous properties of DDPs as effectors in the broader class of polymerase-based DNA writing technologies. The simplicity of CEs should extend their utility to a variety of cells and organisms, and motivate their use to scalably investigate biological questions and systems (e.g. DNA repair mechanisms). More broadly, the modularity of CEs and their potential compatibility with diverse template types should stimulate their continued development and optimization as versatile tools for diverse applications.

## Methods

### Plasmids and oligonucleotides

Plasmid constructs were generated via ligation, isothermal assembly, or Golden Gate assembly. New plasmids generated during this study will be deposited with Addgene (https://www.addgene.org/Benjamin_Kleinstiver/) (**Sup. Table 1**). Target site sequences for gRNAs and pegRNAs are available in **Sup. Table 2**. Expression plasmids for human U6 promoter-driven gRNAs were generated by annealing and ligating duplexed oligonucleotides (oligos) corresponding to spacer sequences into BsmBI-digested BPK1520 (Addgene plasmid 65777)^75^. Expression plasmids for human U6 promoter-driven pegRNAs were generated by phosphorylating, annealing, and ligating 3 sets of duplexed oligos corresponding to (1) the spacer sequence, (2) the SpCas9 gRNA scaffold, and (3) the pegRNA extension (RTT/PBS) into BsmBI-digested MNW320 (see **Sup. Table 1** for additional details regarding the pegRNA cloning protocol). Oligos used in this study for amplicon sequencing (**Sup. Table 3**) and clkDNA oligos (**Sup. Tables 3-6**) were purchased from Integrated DNA Technologies (IDT); gene fragments were ordered from Twist Biosciences.

### Human cell culture and transfection

Human HEK 293T cells (ATCC) were cultured at 37 °C with 5% CO_2_ in Dulbecco’s modified Eagle medium (DMEM) supplemented with 10% heat-inactivated fetal bovine serum and 1% penicillin–streptomycin (ThermoFisher). The supernatant medium from cell cultures was analyzed monthly for the presence of mycoplasma using MycoAlert PLUS (Lonza) or via polymerase chain reaction (PCR).

All experiments were performed with at least 3 replicates; we define biological replicates as results from transfections performed using cells seeded from different passages of cells, and technical replicates as results obtained from transfections performed using the same set of seeded cells.

Transfections were performed between 20-24 hours following seeding of ~2.2×10^4^ HEK 293T cells per well in 96-well plates. Standard transfections included 80 ng of CE expression plasmid, 25 ng of gRNA expression plasmid, 13 ng of nicking gRNA (ngRNA) expression plasmid, and either 16 or 12 pmol of clkDNA unless otherwise indicated (for more details, see **Sup. Note 1**). The DNA mixtures were mixed with TransIT-X2 (Mirus) at a ratio of 0.5 µL of TransIT-X2 per 100 ng of total DNA, in a total volume of 20 µL Opti-MEM (Thermo Fisher Scientific), following manufacturer recommended protocols. This TransIT-X2:DNA solution was mixed gently (very brief low speed vortexing, as aggressive vortexing can negatively impact TransIT-X2:DNA complexing), was incubated for 15 minutes at room temperature, and then gently distributed across the seeded HEK 293T cells, taking care to follow the manufacturer recommendations for preparing the TransIT-X2:DNA complexes (including not leaving the TransIT-X2:DNA complexes in solution for longer than the manufacturer recommended times (e.g. ensuring <30 minutes), pipetting gently to mix the TransIT-X2 and DNA solutions together, and only gently spinning the mixed complexes in a centrifuge for a brief period of time).

### Amplicon sequencing and data analysis

Genomic DNA (gDNA) was harvested ~72 hours after transfection, by discarding the media, resuspending the cells in 100 µL of quick lysis buffer (20 mM Hepes pH 7.5, 100 mM KCl, 5 mM MgCl2, 5% glycerol, 25 mM DTT, 0.1% Triton X-100, and 60 ng/µL Proteinase K (NEB)), heating the lysate for 6 minutes at 66 °C, heating at 98 °C for 2 minutes. Following incubation, gDNA was purified using a 0.8x ratio of paramagnetic beads, prepared as previously described^76,77^. The efficiency of genome modification by CE editing was determined by next-generation sequencing using a 2-step PCR-based Illumina library construction method. Briefly, genomic loci were amplified in a first PCR reaction (PCR-1) reaction from approximately 100 ng of gDNA using Q5 High-fidelity DNA Polymerase (NEB) and the primers listed in **Sup. Table 3**, with cycling conditions of 1 cycle at 98 °C for 2 min; 35 cycles of 98 °C for 10 sec, 65 °C for 10 sec, 72 °C for 20 sec; and 1 cycle of 72 °C for 1 min. PCR-1 products were purified using paramagnetic beads at a ratio of 1.8x. Approximately 20 ng of purified PCR-1 product was used as template for a second PCR (PCR-2) to add Illumina barcodes with adapter sequences using Q5 and the primers listed in **Sup. Table 3**, with cycling conditions of 1 cycle at 98 °C for 2 min; 10 cycles at 98 °C for 10 sec, 65 °C for 30 sec, 72 °C 30 sec; and 1 cycle at 72 °C for 5 min. PCR-2 products were pooled based on concentrations from capillary electrophoresis (QIAxcel, Qiagen). Final libraries were quantified by Qubit dsDNA High Sensitivity assay (ThermoFisher) and sequenced on a MiSeq sequencer using a 300-cycle v2 kit (Illumina). On-target genome editing activities were determined from amplicon sequencing data using CRISPResso2 (ref. ^78^).

Using CRISPResso2, amplicon sequences were aligned to a reference sequence in HDR mode using the intended editing outcome as the expected allele (-e) and the parameters “-q 30” and “-discard_indel_reads”. For each amplicon, the quantification window (-qwc) was defined as the entire sequence between gRNA- and ngRNA-directed cut sites plus an additional 10 bp on either side of each nicking site. The same quantification window was used to analyze data for each amplicon, whether or not a ngRNA was transfected. Editing efficiencies were quantified by determining: (# of reads aligned to HDR / number of total reads). Indel efficiencies were quantified as (number of discarded indel-containing reads / number of total reads). Analysis of experiments containing clkDNAs with repair-evading substitutions was run using CRISPResso2 in standard mode, providing the reference amplicon and the gRNA sequence, and using the same quantification window as in HDR mode.

Reads containing template insertions (clkDNA or pegRNA scaffold) were analyzed using BBduk from the bbtools suite^79^, reads were merged and filtered for Q>30, minlen=100. To determine template insertion, merged and filtered FastQ files were imported into Geneious Prime (v2022.1.1) and manually searched. Reads containing the gRNA scaffold sequence and/or the PCV2 recognition sequence as well as reads containing the correct edit were counted. Scaffold integration was calculated as the (number of reads containing scaffold or PCV2 sequence) / (total number of reads containing the intended edit)*100.

### Assessment of gRNA-dependent off-targets

We examined putative gRNA-dependent off-target editing with CEs by first designing 12 off-target sites per primary gRNA using CasOFFinder^63^ with search parameters of 1-3 mismatches, 20 nt spacer, NRG protospacer-adjacent motif (PAM), and no RNA or DNA bulge off-targets. When possible, to increase potential sensitivity to detect off-targets, the CasOFFinder output was currated to select off-target sites with minimal mismatches in seed region (~10 bp adjacent to PAM). Amplicon-specific primers to amplify off-target sites were designed using Primer3 with amplicon length between 150-250 bp, and off-target sequence dictated as the target region and ideal melting temperature between 63 and 68°C. The PCR-1 amplicon-specific primers were designed by adding Illumina adapter sequences to gene-specific sequences. Transfections for CEs were performed as described above; nuclease transfections contained 80 ng of SpCas9 nuclease or nickase expression plasmid and 25 ng of the primary gRNA. gDNA was harvested, amplicon PCRs were performed, sequencing libraries were prepared and sequenced as described above. For analysis, CRISPResso2 was run in standard mode and indels were calculated using the CRISPResso_quantification_of_editing_frequency.txt output as (SUM(Insertions, Deletions)-Insertions and Deletions)/(Reads_Total)*100. On target editing for the *DNMT1*, *ACTB*, and *VEGFA* targets was analyzed using CRISPREsso2 HDR mode.

### Assessment of potential HUHe off-targets

HUH pseudo-site targets were designed using TagScan^80^ to search for sites within the human reference genome with the PCV2 binding sequence (AAGTATTACCAGC) within 20 bp of an NGG PAM, optimally placing the PCV2 binding site in the solvent-accessible PAM-distal region of the non-target strand (**Sup. Table 7**). Once putative target sites were identified, oligonucleotides were ordered for spacer sequences and gRNA plasmids were cloned as described above. Transfections were performed as described above, containing 80 ng of enzyme expression plasmid (CE, CE-deadPCV2, CE-deadCas9, SpCas9(H840A), or SpCas9), 25 ng of HUH pseudo-site targeting gRNA plasmid, optionally, 12 pmol of clkDNA, and 0.5 µL/100 ng of TransIT-X2 were mixed into a total volume 20 µL of Opti-MEM, and transfections, gDNA preparation, and sequencing protocols were performed as described above.

## Data availability

Primary datasets will be made available in **Supplementary Tables**. Sequencing datasets will be deposited with the NCBI Sequence Read Archive (SRA) under <u>BioProject ID PRJNA1015647</u>.

## Acknowledgements

We acknowledge members of the Kleinstiver laboratory (for critical feedback), A. Anzalone (for advice regarding data analysis), E.J. Sontheimer and W. Xue for communicating results prior to publication, M.N. Whittaker for cloning the pegRNA entry vector, and Z. Hebert and M. Berkeley (DFCI MBCF for sequencing support). J.F.daS was supported by an EMBO Long Term Fellowship (ALTF 750-2022), C.J.T. was supported by a National Science Foundation Graduate Research Fellowship (2020295403), L.M. was supported by a Massachusetts General Hospital (MGH) Executive Committee on Research (ECOR) Fund for Medical Discovery Fundamental Research Fellowship Award, and research in the B.P.K lab was supported by an MGH Howard M. Goodman Fellowship, an MGH Research Scholar Award 2023-2028, and National Institutes of Health (NIH) grant DP2-CA281401.

## Author contributions

C.J.T. conceived of initial concepts. J.F.daS, C.J.T., and B.P.K. contributed to subsequent ideation. J.F.daS, C.J.T., M.L.E., E.M.K., L.M., and D.R.R. designed and performed experiments. B.P.K. contributed to experimental design and oversaw the study. J.F.daS., C.J.T., and B.P.K. wrote the manuscript with contributions and/or revisions from all authors.

## Competing interests

J.F.daS, C.J.T, and B.P.K. are inventors on a patent application filed by Mass General Brigham (MGB) that describes click editing. C.J.T. and B.P.K. are inventors on additional patents or patent applications filed by MGB that describe genome engineering technologies. B.P.K. is a consultant for EcoR1 capital and is on the scientific advisory board of Acrigen Biosciences, Life Edit Therapeutics, and Prime Medicine. B.P.K. has a financial interest in Prime Medicine, Inc., a company developing therapeutic CRISPR-Cas technologies for gene editing. B.P.K.’s interests were reviewed and are managed by MGH and MGB in accordance with their conflict-of-interest policies. The other authors declare no competing interests.

