## Supplementary Notes and Figures for "Click editing enables programmable genome writing using DNA polymerases and HUH endonucleases"

### Supplementary Materials

Supplementary Tables

**Supplementary Table 1:** Plasmids

**Supplementary Table 2:** gRNA target sites

**Supplementary Table 3:** Amplicon sequencing primers

**Supplementary Table 4:** clkDNA oligonucleotides

**Supplementary Table 5:** clkDNA oligonucleotides for clkDNA screens

**Supplementary Table 6:** clkDNA oligonucleotides with additional mismatches

**Supplementary Table 7:** HUH pseudosites

Supplementary Notes

**Supplementary Note 1:** Optimization of clkDNA dose p.2

**Supplementary Note 2:** Analysis of click editor controls and indels p.2

**Supplementary Note 3:** Expanded discussion of HUHe off-target analysis p.3

Supplementary Figures and Legends

**Supplementary Figures 1 - 15** p.5

Supplementary References p.23

#### Supplementary Notes:

##### Supplementary Note 1. Optimization of clkDNA dose.

To determine the optimal dose of clkDNA to use for click editing experiments, we co-transfected HEK 293T cells with CE1 (PCV2-nSpCas9(H840A)-EcKlenow), a primary gRNA targeting *DNMT1*, and a secondary ngRNA (n2(+49)), together with increasing doses of a clkDNA (from 0-32 pmols) encoding a +3-5 AGG deletion (**Sup. Fig. 2a**). In a control condition, we co-transfected an nSpCas9(H840A) with both gRNAs and a similar clkDNA lacking the PCV2 recognition site (i.e. only PT and PBS) (**Sup. Fig. 2a**). Our results confirmed that productive click editing only occurs in the CE condition. Moreover, the optimal clkDNA dose for this experiment ranged between 10 and 16 pmols which, considering the length and composition of this particular oligonucleotide, is equivalent to 128.8-206.1 ng of ssDNA. The highest dose tested (32 pmols) induced some toxicity to the cells, which might explain the decrease in click editing efficiency. Considering these results, for a selection of subsequent experiments (**Fig.1 f-l; Sup. Fig.3; Sup. Figs. 4a-d**) we used 16 pmols of clkDNA and 0.6 μL per 100 ng of total DNA of transfection reagent (TransIT-X2, Mirus). However, upon repeating transfections several times, we noticed that in some cases these conditions could induce toxicity, a potential cofounding factor for accurately determining editing efficiencies. We also hypothesized that high doses of clkDNA may decrease editing efficiencies through spacer sequestration (due to sequence complementarity[1]), or RNaseH-mediated degradation of DNA:RNA hybrid created by pairing in solution between a clkDNA and the spacer of a gRNA. We then performed an additional test to determine whether decreasing the clkDNA dose to 12 pmols would improve editing and viability (**Sup. Fig 2b**). In this experiment, we also decreased the concentration of transfection reagent to 0.5 μL per 100 ng of total DNA. These new conditions led to a 1.37-fold increase in click editing efficiency compared to the previous experimental setup with no alteration in indel levels (**Sup. Fig 2b**), and significantly less general toxicity to cells. All experiments for the remainder of our study were performed with these optimized conditions (12 pmols of clkDNA and 0.5 μL of TransIT-x2 per 100 ng of total transfected DNA – in **Figs. 2, 3** and **4**; and **Sup. Figs. 4e** and **5-15**). We envision that further optimization of clkDNA chemistry, as well as alternative delivery modalities (e.g. RNPs), will permit a reduction of clkDNA dosage to alleviate any clkDNA:spacer sequestration of RNaseH-mediated gRNA cleavage without compromising editing efficiencies.

##### Supplementary Note 2. Analysis of click editor controls and indels.

We achieved productive and precise click editing when using our original CE construct (PCV2-nCas9(H840A)-EcKlenow; **Figs. 1e**-**g**). To confirm that all components were essential for productive click editing, we performed experiments using a series of control conditions that included: (1) a nickase Cas9 (H840A) (nCas9) with a clkDNA that did not contain an HUH site (i.e. PBS/PT only); (2) a CE containing a dead Cas9 (H840A, D10A) but a functional PCV2 and EcKlenow; (3) a CE containing a dead PCV2 (Y96F) (dPCV2)[2] but a fully functional nCas9 and EcKlenow; and (4) a CE containing an attenuated EcKlenow (D355A, D357A) (dKlenow)[3,4] but a fully functional nCas9 and PCV2 (**Figs 1f**,**g**). Although our control conditions confirmed the essentiality of all active components for productive click editing, we could still detect low level precise editing when using a CE containing a dPCV2 or dKlenow. For click editing with a dPCV2-CE, it is possible that the catalytically inactivated PCV2 domain may retain some residual DNA binding affinity (but be unable to catalyze covalent adduct formation with a clkDNA), enabling weak localization of the clkDNA to the target site-bound dPCV2-CE (which contains a fused, fully active EcKlenow). Alternatively, the PBS from a clkDNA from solution could hybridize with the nCas9-induced NTS flap, providing weak but sufficient annealing of the clkDNA:NTS duplex to initiate polymerization. When testing the CE construct fused to dKlenow, a clkDNA should be covalently localized to the target site via PCV2 but polymerization should be substantially attenuated from the fused polymerase[4] (or any other CE-fused polymerase in trans). In this case, the low levels of precise click editing that we observed may be the result of incomplete inactivation or EcKlenow, or endogenous polymerases interacting with the clkDNA-NTS hybrid to initiate polymerization.

In this experiment we also compared CE1 with CE1.n2 and CE1.n2b (**Figs 1f**,**g**). While CE1.n2 employs a secondary gRNA to direct CE-mediated nicking (ngRNA) against the non-edited strand at a certain distance upstream or downstream of the primary nick site, CE1.n2b uses a ngRNA to nick the non-edited strand only after the edit is installed (**Sup. Fig. 1**). Because the CE1.n2b strategy minimizes concurrent nicks on opposite strands, the probability of the reaction resulting in insertion or deletion mutations (indels) is reduced (similarly to as observed with PE3 and PE3b nicking events for prime editing[5]) (**Fig. 1g**).

**Supplementary Note 3: Expanded discussion of HUHe off-target analysis.**

Aside from their use to tether ssDNA templates as homology-directed repair donors to Cas9-induced sites of DNA breaks[6], the uses of HUHes have been largely unexplored in the context of genome editing experiments. We sought to better understand any potential impacts of overexpressing an HUHe in human cells, which we imagined might be relatively innocuous given HUHes dual modes of specificity: (1) being specific for ssDNA (and not dsDNA) and (2) requiring a specific DNA binding motif on the substrate[7]. While the vast majority of the human genome is double-stranded in most cell types, regions of ssDNA can become transiently exposed during transcription and/or DNA replication, including genomic hotspots that are prone to ssDNA deamination events from endogenous deaminases[8–10]. Fortunately, RPA, hSSB, and other process-specific factors can occupy these regions to limit access or damage to ssDNA. Moreover, these ssDNA regions tend to be transient; thus, sustained solvent exposure of a specific region of ssDNA containing the full HUHe recognition sequence is unlikely. Thus, we wondered whether intentionally overexpressed HUHes could act on perfectly or partially matched HUHe binding pseudosites, which may exist in the human genome during click editing.

In an initial experiment, we identified genomic sites bearing a perfectly matched PCV2 HUHe binding sequence (AAGTATTACCAGC) using TagScan[11]. After constitutively overexpressing a CE or HUHe domain in cells for ~72 hours, we extracted genomic DNA and sequenced these regions to determine whether we observed an increase in indels in HUHe-treated conditions. Despite performing our experiments in rapidly dividing and highly transcriptionally active HEK 293T cells, we were unable to detect indels at these genomic HUHe sites at levels above untransfected controls (**Figs. 4j-l** and **Sup. Fig. 14d**,**e**).

To improve our sensitivity to detect potential HUHe-genomic interactions, we searched for and nominated additional PCV2-HUHe sites within 20 bp of an NGG PAM, optimally placing the PCV2 binding site in the solvent-accessible PAM-distal region of the non-target strand (to improve the sensitivity of potentially detecting HUHe-based indels by artificially maximizing access of the CE or HUHe to the ssDNA NTS).

To test this possibility, we performed a reporter experiment where we intentionally targeted the CE (with nCas9 or dCas9) to HUHe genomic sites to induce an R-loop (**Sup. Fig. 14a**). By artificially creating stable R-loops, we were able to detect low level indels at levels slightly greater than control conditions (**Sup Figs. 14b**,**c**). We observed indels only when the HUHe pseudosite was in a very specific region of the Cas9 target site, an effect that was dampened slightly by the addition of a clkDNA in the transfections. We anticipate that stable R-loops at these very sparse/rare genomic locations containing an HUHe recognition sequences should occur infrequently if at all, though future studies are required to more deeply interrogate the genome-scale impact of HUHe overexpression.

#### Supplementary Figures and Legends

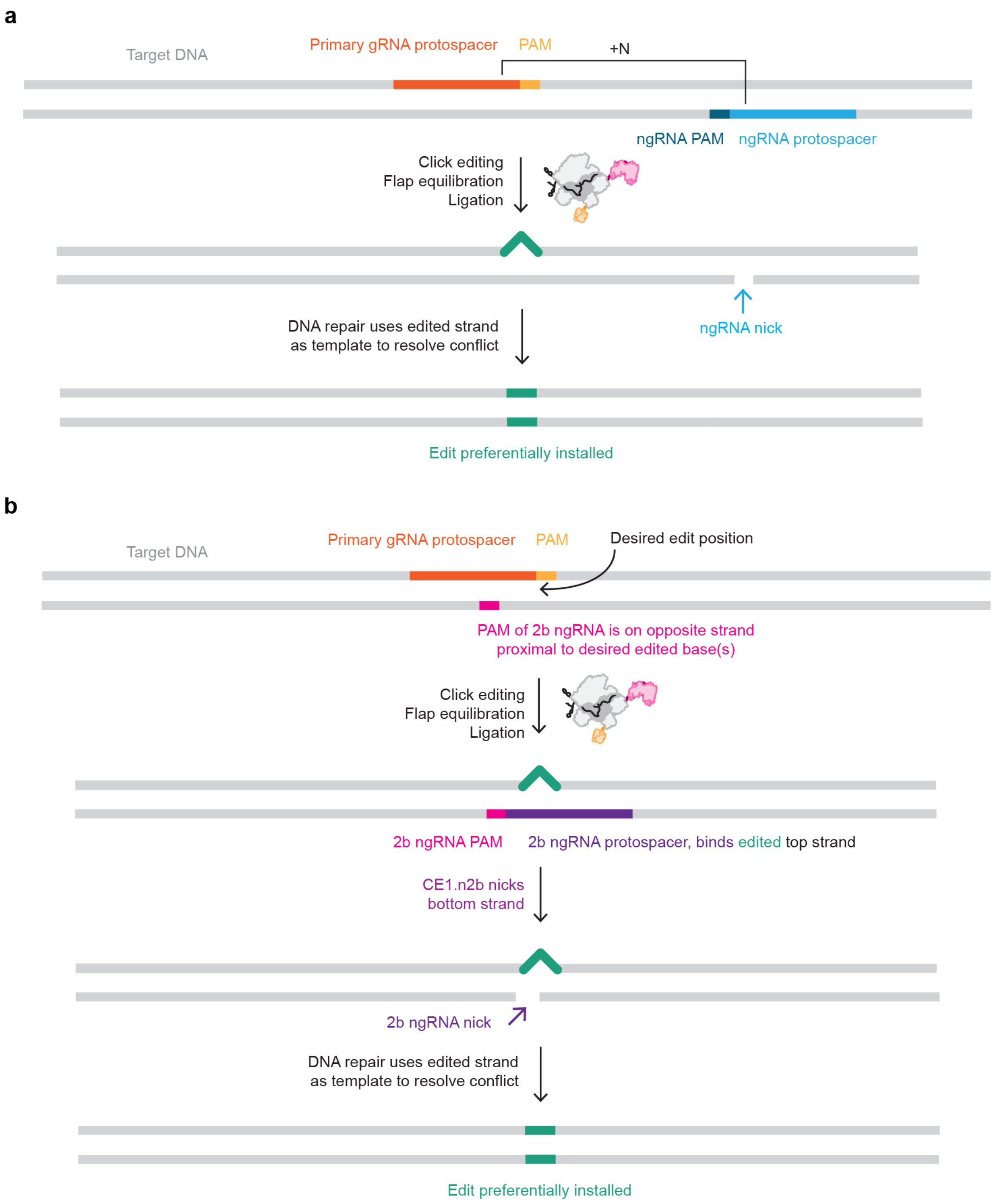


**Supplementary Figure 1. Schematic of CE1.n2 and CE1.n2b compositions**. **a,b,** The use of a secondary nicking gRNA (ngRNA) for CE1.n2 or CE1.n2b conditions (when the ngRNA is located distal from the edit or overlaps the edit as shown in **panels a** and **b**, respectively) can modify click editing efficiency and/or the level of insertion or deletion mutations (indels) observed. The n2 and n2b nicking conventions are similar to the PE3 and PE3b nicking approaches for PEs[5].

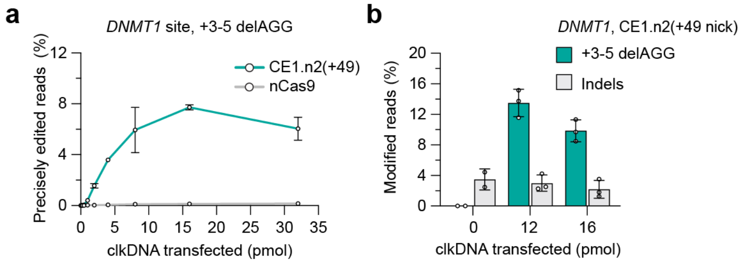


**Supplementary Figure 2. Impact of clkDNA concentration on click editing efficiency**. **a**, Titration of clkDNA dose (0 - 32 pmols) for installing a +3-5 AGG deletion in the *DNMT1* locus using CE1.n2 or only nSpCas9(H840A). **b**, Comparison of precise editing efficiency and insertion or deletion mutations (indels) for the *DNMT1* site using CE1.n2 with 12 or 16 pmols of clkDNA. Data in **panels a** and **b** from HEK 293T cell experiments; mean, s.d., and individual datapoints shown for n=3 independent technical replicates.

a

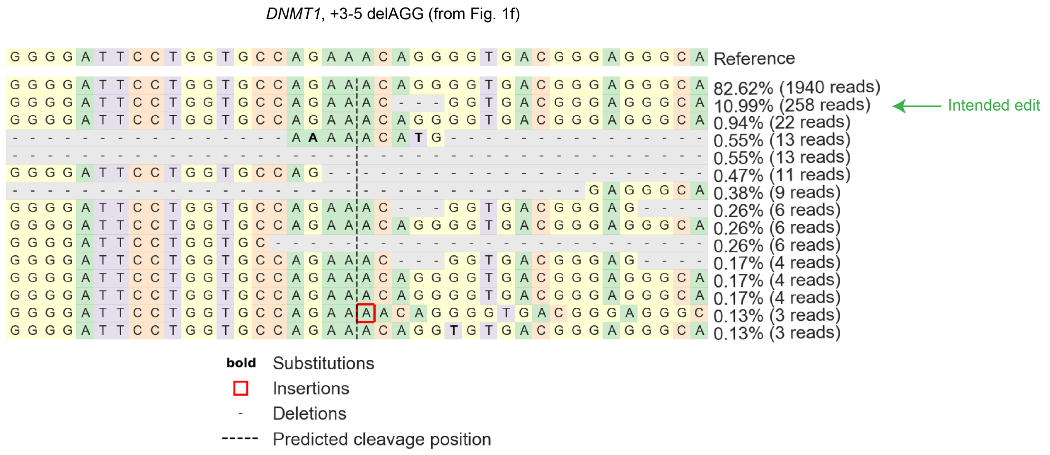


b

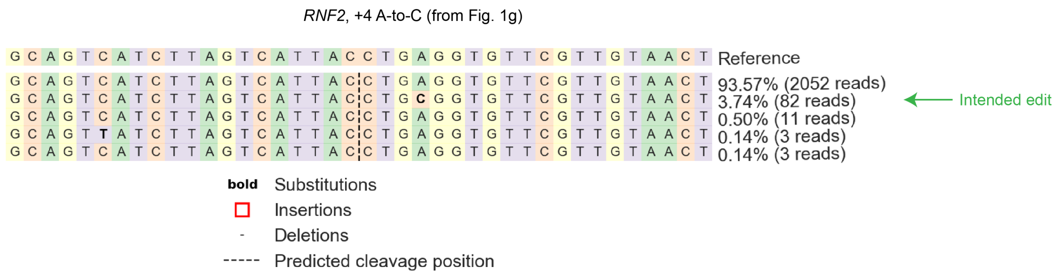


**Supplementary Figure 3. Edit outcomes from initial click editing experiments**. **a-b**, Representative allele frequency tables from CRISPResso2 (ref. [12]) output for installation of a +3-5 AGG deletion in the *DNMT1* locus (with CE1.n2(+49)) or a +4 A-to-C transversion in the *RNF2* locus (with CE1.n2b(+4)) (**panels a** and **b**, respectively).

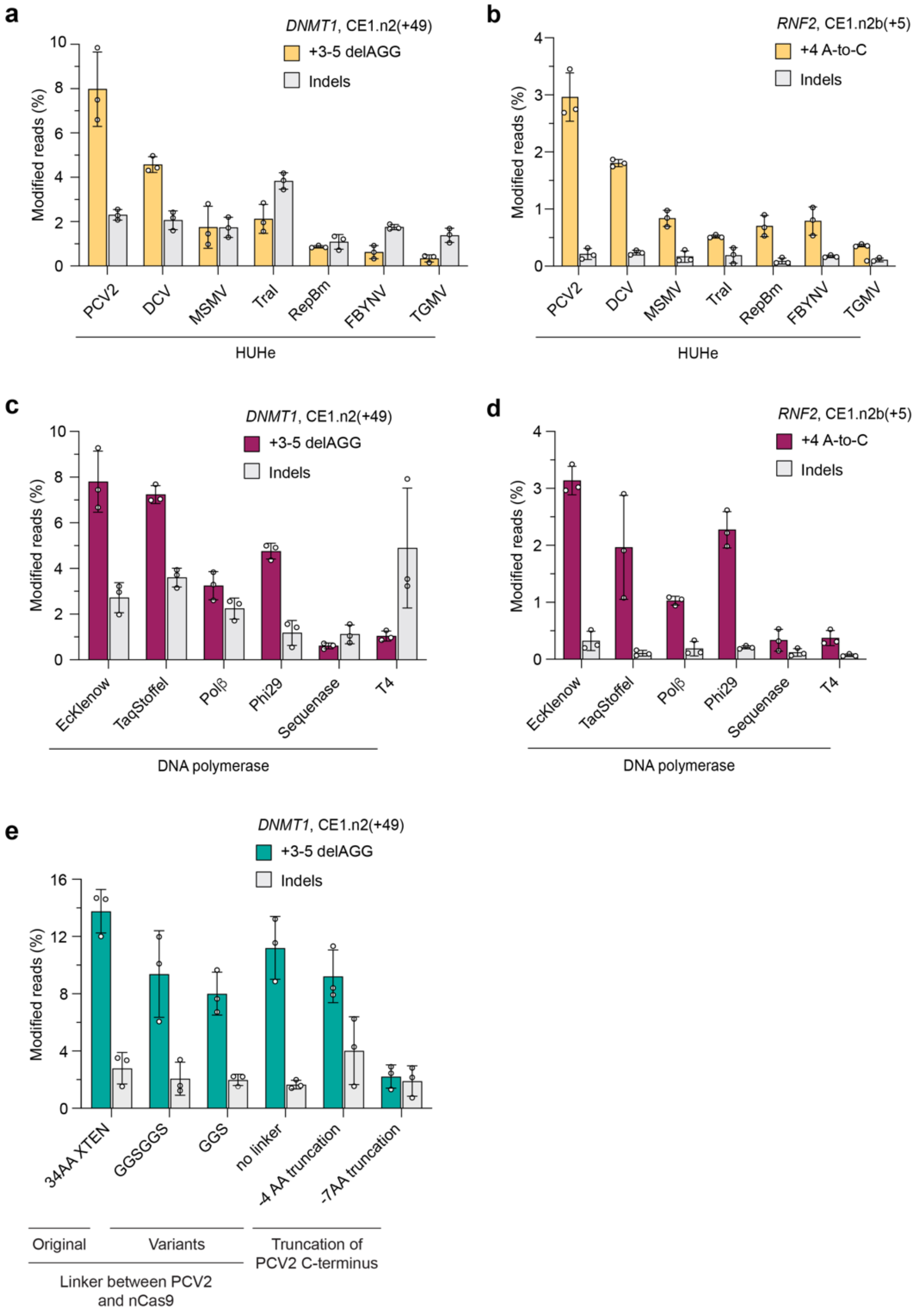


**Supplementary Figure 4. Assessment of different click editor architectures. a-b**, Percentage of sequencing reads with precise edits or insertion or deletion mutations (indels) when using CE constructs encoding different HUHe domains to install edits at *DNMT1* or *RNF2* (**panels a** and **b**, respectively). PCV2, porcine circovirus 2; DCV, duck circovirus; MSMV, maize striate mosaic virus; TraI, *E.coli* conjugation protein TraI*;* RepBm, RepB *Fructobacillus tropaeola*; FBYNV, fava bean necrosis yellow virus; TGMV, tomato golden mosaic virus. **c-d,** Percentage of sequencing reads with precise edits or indels when using CE constructs encoding different DNA-dependent DNA polymerases installing edits at *DNMT1* or *RNF2* (**panels c** and **d**, respectively). TaqStoffel, Stoffel fragment from *Thermus aquaticus* DNA polymerase; Polβ, human polymerase beta; Phi29, DNA polymerase from bacteriophage ϕ29 (D169A); Sequenase, engineered truncation of T7 bacteriophage DNA polymerase; T4, T4 bacteriophage DNA polymerase. **e**, Percentage of sequencing reads with precise edits or indels at the *DNMT1* locus using CE constructs encoding different linker variants between PCV2 and nSpCas9(H840A), as well as different PCV2 C-terminal truncations. Data in **panels a-e** from HEK 293T cell experiments; mean, s.d., and individual datapoints shown for n=3 independent technical replicates.

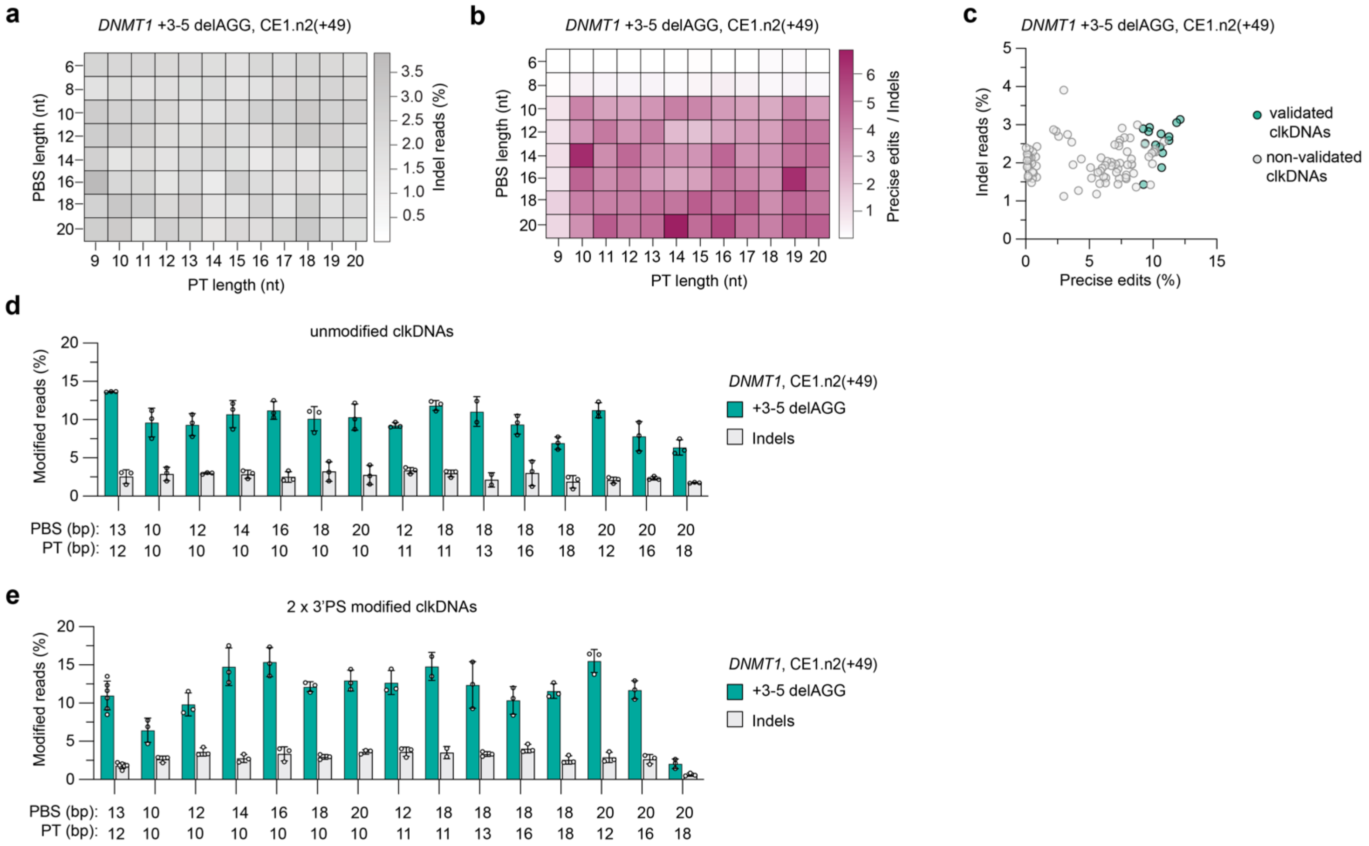


**Supplementary Figure 5. *DNMT1* clkDNA screens and validation. a**, Percentage of sequencing reads harboring indels at the *DNMT1* locus, using CE1.n2 and clkDNAs with varying PBS and PT lengths to install a +3-5 AGG deletion. **b**, Ratio of precise editing to indels at the *DNMT1* locus, using CE1.n2 and clkDNAs with varying PBS and PT lengths. **c**, Scatter plot depicting percentage of edit and indels at the *DNMT1* locus for CE1.n2 and clkDNAs with varying PS and PT lengths. Highlighted are the clkDNAs that led to some of the highest levels of editing, which we selected for validation. **d,e,** Percentage of sequencing reads with precise edits or indels at the *DNMT1* locus with selected clkDNAs which are either unprotected or 2x3'PS protected (**panels d** and **e**, respectively), using CE1.n2. Data in **panels a-c** from HEK 293T cell experiments; mean, s.d., and individual datapoints shown for n = 3 independent biological replicates. Data in **panels d** and **e** from HEK 293T cell experiments; mean, s.d., and individual datapoints shown for n = 3 independent technical replicates.

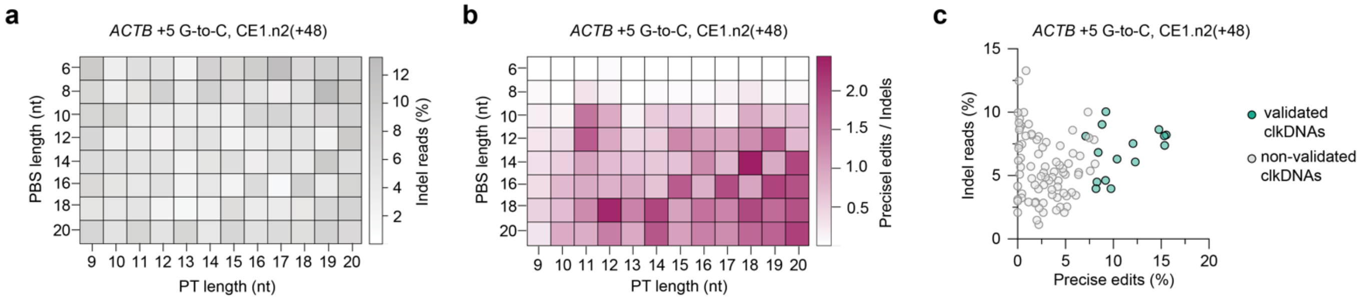


**Supplementary Figure 6. *ACTB* clkDNA screens and validation. a,** Percentage of sequencing reads harboring indels at the *ACTB* locus, using CE1.n2 and clkDNAs with varying PBS and PT lengths to install a +5 G-to-C substitution. **b**, Ratio of precise editing to indels at the *ACTB* locus, using CE1.n2 and clkDNAs with varying PBS and PT lengths. **c**, Scatter plot depicting percentage of edit and indels at the *ACTB* locus for CE1.n2 and clkDNAs with varying PS and PT lengths. Highlighted are the clkDNAs that led to some of the highest levels of editing, which we selected for validation. Data in **panels a-c** from HEK 293T cell experiments; mean, s.d., and individual datapoints shown for n = 3 independent biological replicates.

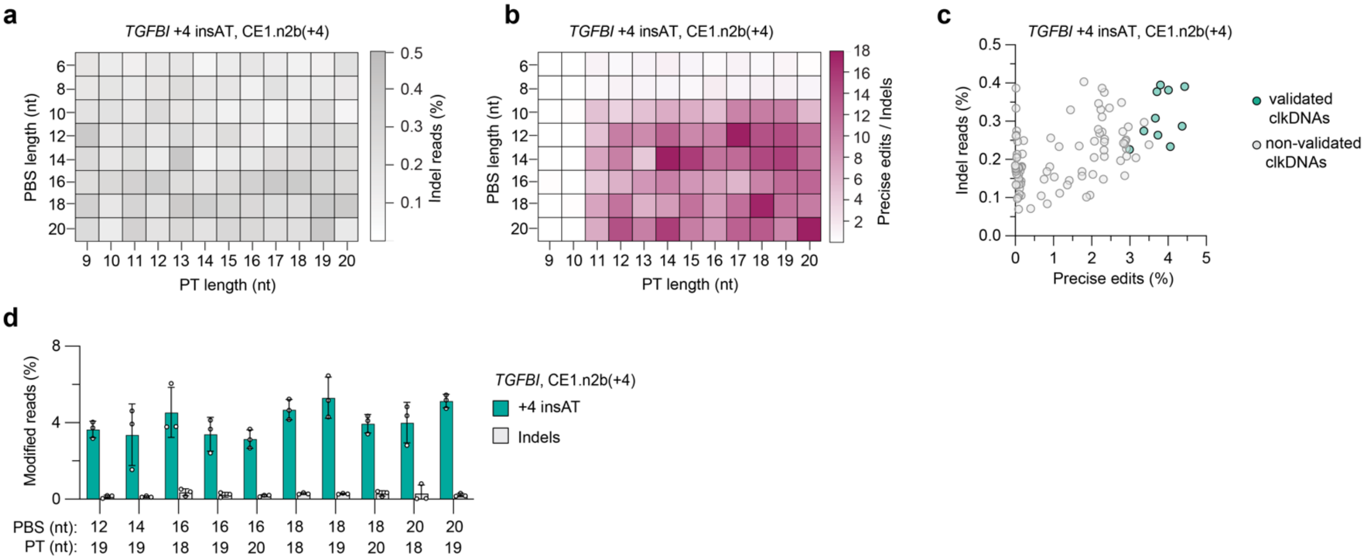


**Supplementary Figure 7. *TGFBI* clkDNA screens and validation**. **a,** Percentage of sequencing reads harboring indels at the *TGFBI* locus, using CE1.n2 and clkDNAs with varying PBS and PT lengths to install a +4 AT insertion. **b**, Ratio of precise editing to indels at the *TGFBI* locus, using CE1.n2 and clkDNAs with varying PBS and PT lengths. **c**, Scatter plot depicting percentage of edit and indels at the *TGFBI* locus for CE1.n2 and clkDNAs with varying PS and PT lengths. Highlighted are the clkDNAs that led to some of the highest levels of editing, which we selected for validation. **d,** Percentage of sequencing reads with precise edits or indels at the *TGFBI* locus with selected clkDNAs which are 2x3'PS protected, using CE1.n2. Data in **panels a-c** from HEK 293T cell experiments; mean, s.d., and individual datapoints shown for n = 3 independent biological replicates. Data in **panel d** from HEK 293T cell experiments; mean, s.d., and individual datapoints shown for n = 3 independent technical replicates.

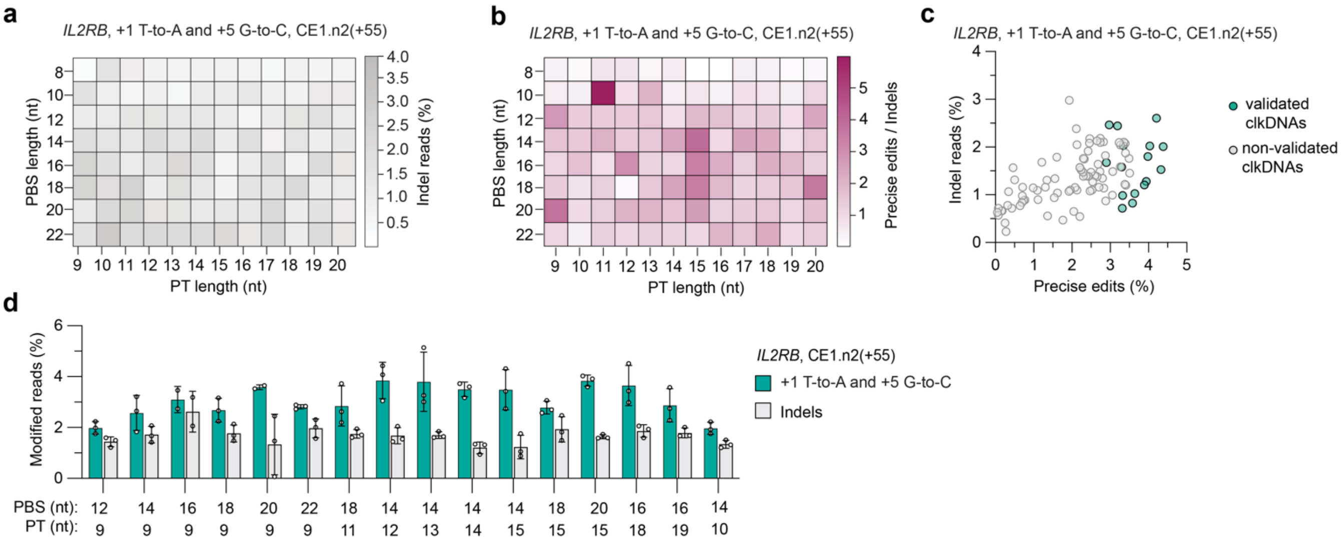


**Supplementary Figure 8. *IL2RB* clkDNA screens and validation**. **a**, Percentage of sequencing reads harboring indels at the *IL2RB* locus, using CE1.n2 and clkDNAs with varying PBS and PT lengths to install a dual +1 T-to-A and +5 G-to-C edit. **b**, Ratio of precise editing to indels at the *IL2RB* locus, using CE1.n2 and clkDNAs with varying PBS and PT lengths. **c**, Scatter plot depicting percentage of edit and indels at the *IL2RB* locus for CE1.n2 and clkDNAs with varying PS and PT lengths. Highlighted are the clkDNAs that led to some of the highest levels of editing, which we selected for validation. **d,** Percentage of sequencing reads with precise edits or indels at the *IL2RB* locus with selected clkDNAs which are 2x3'PS protected, using CE1.n2. Data in **panels a-c** from HEK 293T cell experiments; mean, s.d., and individual datapoints shown for n = 3 independent biological replicates. Data in **panel d** from HEK 293T cell experiments; mean, s.d., and individual datapoints shown for n = 3 independent technical replicates.

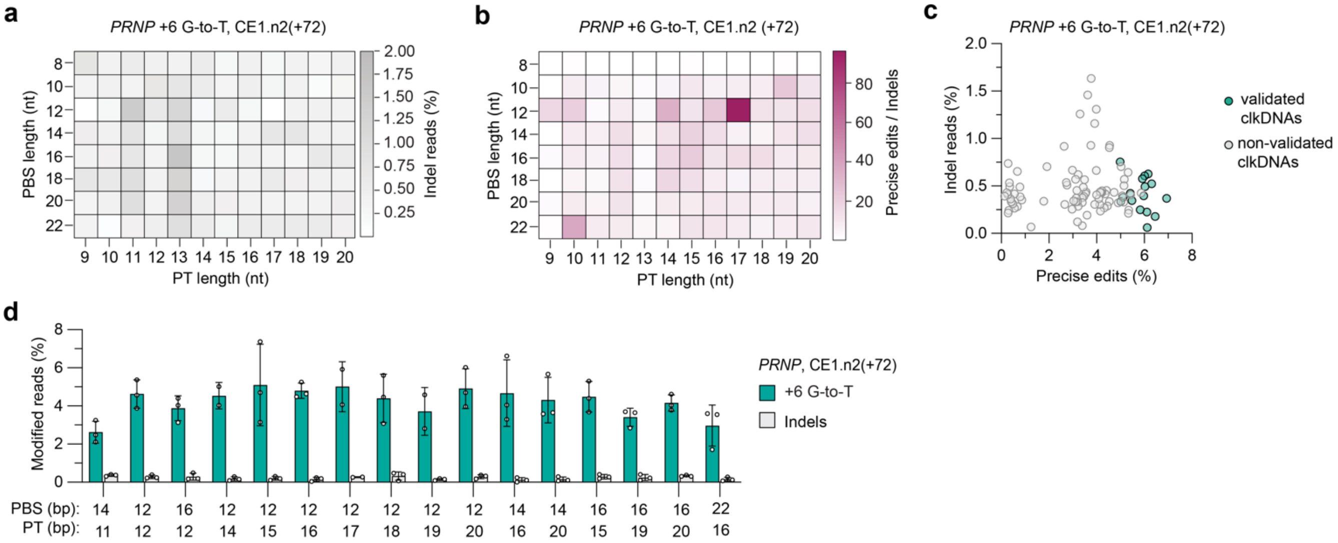


**Supplementary Figure 9. *PRNP* clkDNA screens and validation**. **a,** Percentage of sequencing reads harboring indels at the *PRNP* locus, using CE1.n2 and clkDNAs with varying PBS and PT lengths to install a +6 G-to-T edit. **b**, Ratio of precise editing to indels at the *PRNP* locus, using CE1.n2 and clkDNAs with varying PBS and PT lengths. **c**, Scatter plot depicting percentage of edit and indels at the *PRNP* locus for CE1.n2 and clkDNAs with varying PS and PT lengths. Highlighted are the clkDNAs that led to some of the highest levels of editing, which we selected for validation. **d,** Percentage of sequencing reads with precise edits or indels at the *PRNP* locus with selected clkDNAs which are 2x3'PS protected, using CE1.n2. Data in **panels a-c** from HEK 293T cell experiments; mean, s.d., and individual datapoints shown for n = 3 independent biological replicates. Data in **panel d** from HEK 293T cell experiments; mean, s.d., and individual datapoints shown for n = 3 independent technical replicates.

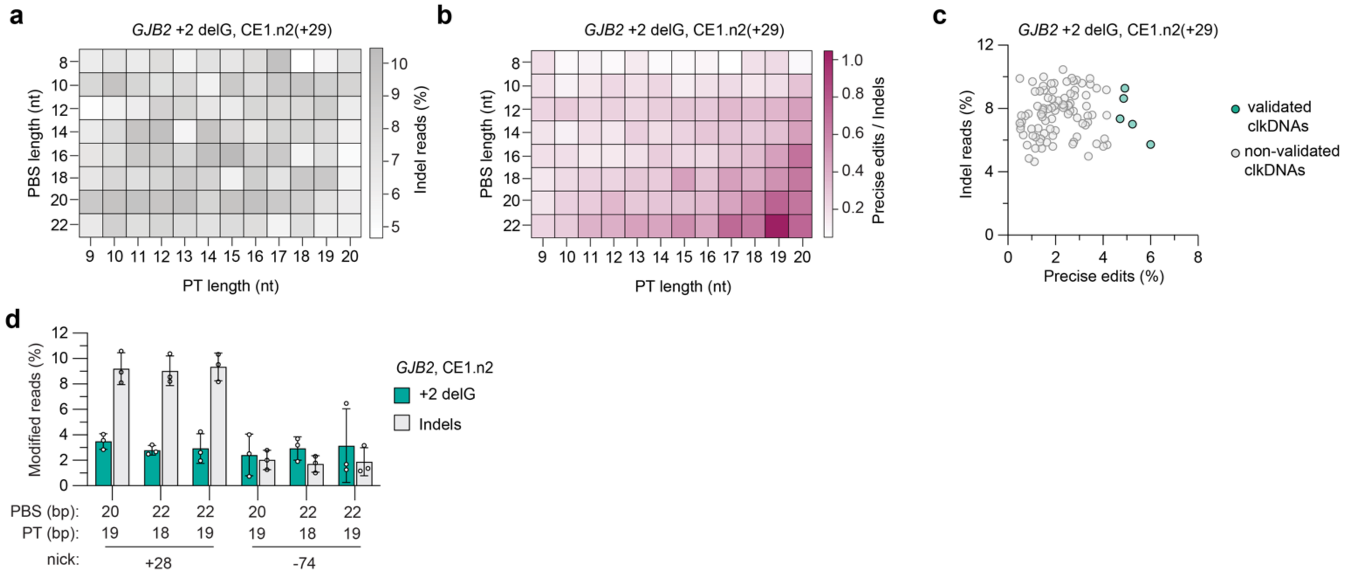


**Supplementary Figure 10. *GJB2* clkDNA screens and validation. a,** Percentage of sequencing reads harboring indels at the *GJB2* locus, using CE1.n2 and clkDNAs with varying PBS and PT lengths to install a +2 G deletion. **b**, Ratio of precise editing to indels at the *GJB2* locus, using CE1.n2 and clkDNAs with varying PBS and PT lengths. **c**, Scatter plot depicting percentage of edit and indels at the *GJB2* locus for CE1.n2 and clkDNAs with varying PS and PT lengths. Highlighted are the clkDNAs that led to some of the highest levels of editing, which we selected for validation. **d,** Percentage of sequencing reads with precise edits or indels at the *GJB2* locus with selected clkDNAs which are 2x3'PS protected, using CE1.n2 and two different ngRNAs. Data in **panels a-c** from HEK 293T cell experiments; mean, s.d., and individual datapoints shown for n = 3 independent biological replicates. Data in **panel d** from HEK 293T cell experiments; mean, s.d., and individual datapoints shown for n = 3 independent technical replicates.

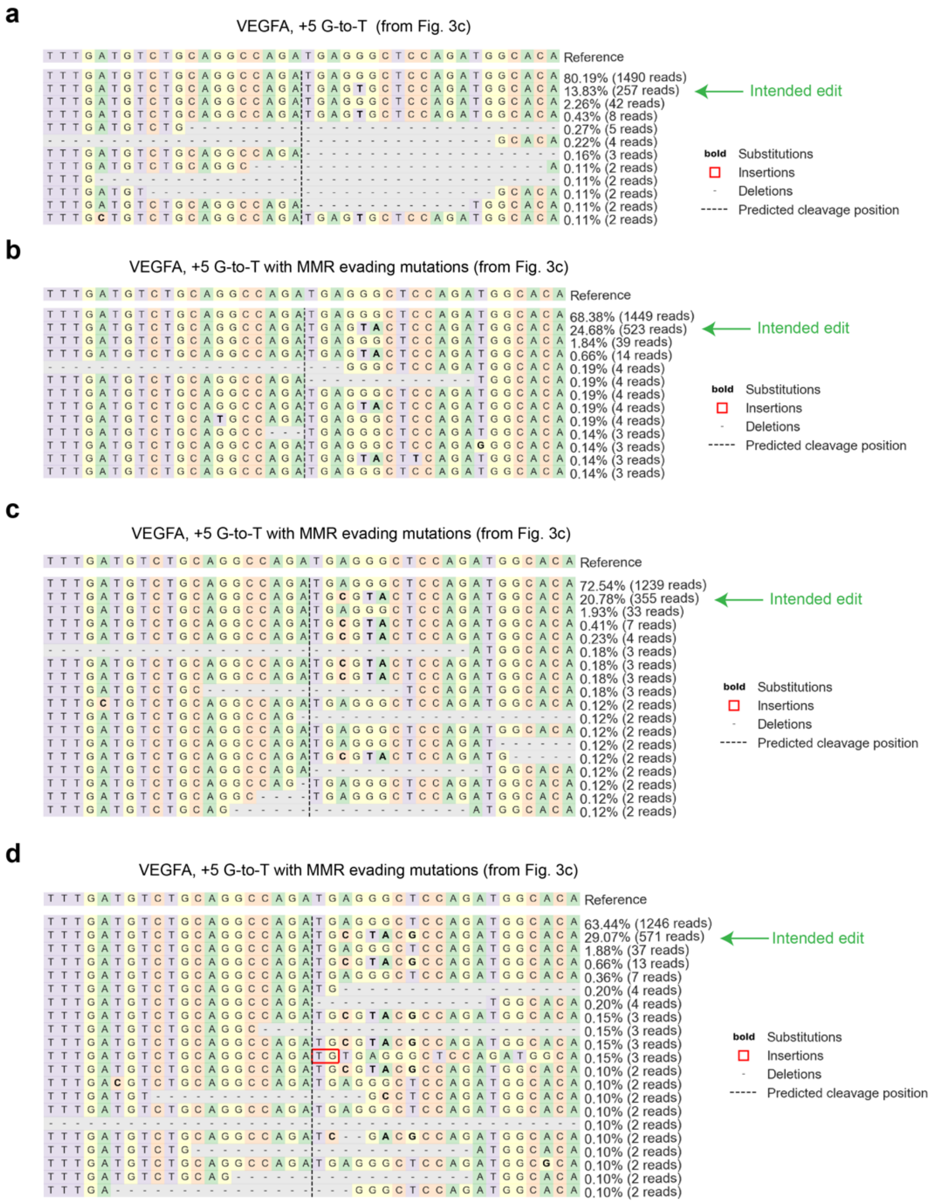


**Supplementary Figure 11. Exemplary allele outcome tables from CE experiments using various clkDNAs. a-d,** Representative allele frequency tables from CRISPResso2 (ref. [12]) outputs for the installation of a +5 G-to-T transversion at the *VEGFA* locus using a clkDNA with only the intended edit (**panel a**), or when using other clkDNAs encoding additional substitutions to evade MMR (**panels b-d**).

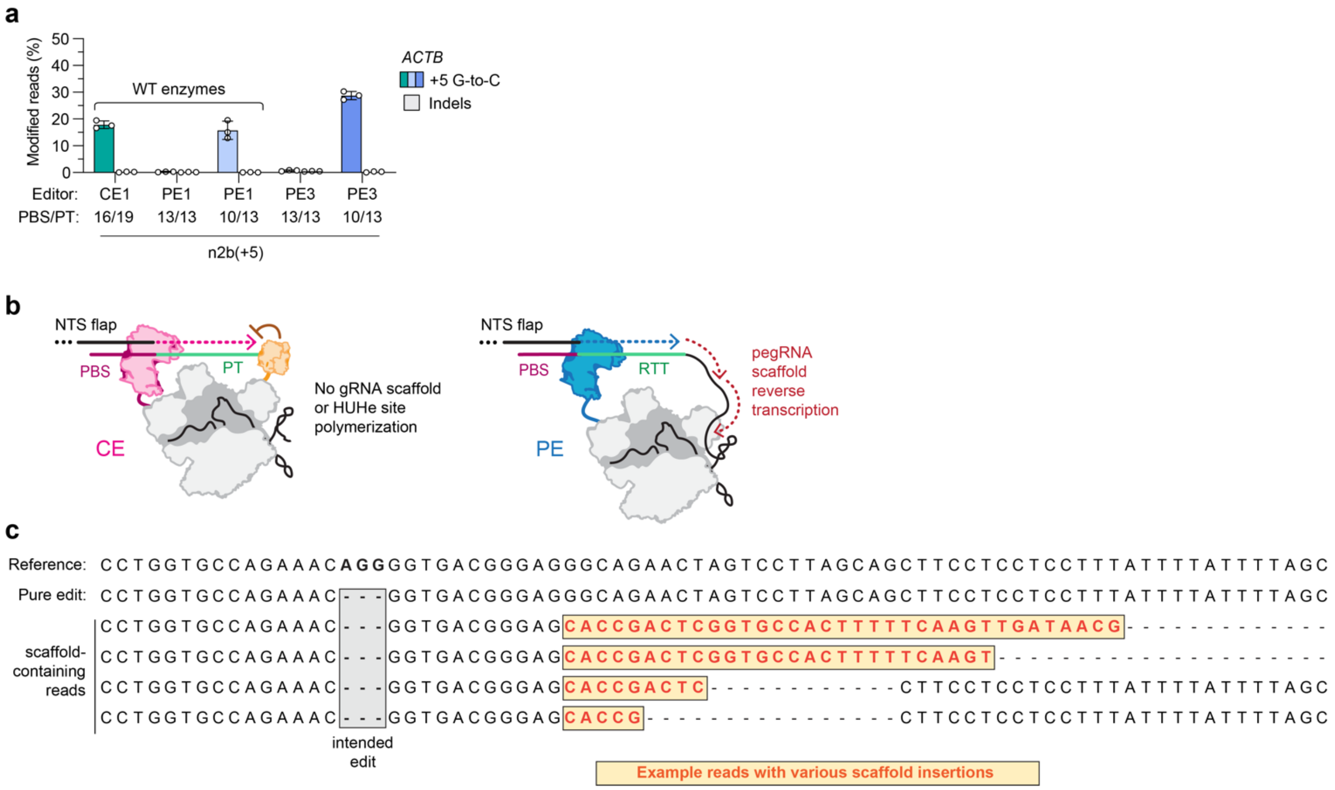


**Supplementary Figure 12. Comparison of unwanted template-mediated insertions with CEs and PEs**. **a,** Percentage of sequencing reads with precise edits or indels at the *ACTB* locus with CE1, PE1, or PE3, with a +5 n2b gRNA. Data from HEK 293T cell experiments; mean shown for n = 3 independent biological replicates. WT, wild-type. **a**, Schematic of templated polymerization for CEs and PEs. With CEs, the HUHe site on the clkDNA is blocked by the bound HUHe to prevent read-though. During prime editing experiments, RT-mediated read-through into the pegRNA scaffold sequence can lead to unwanted insertion byproducts[5]. **c,** Example reads of scaffold insertion products when using PE2 to install a 3-bp deletion at *DNMT1*, analyzed as described in the Methods.

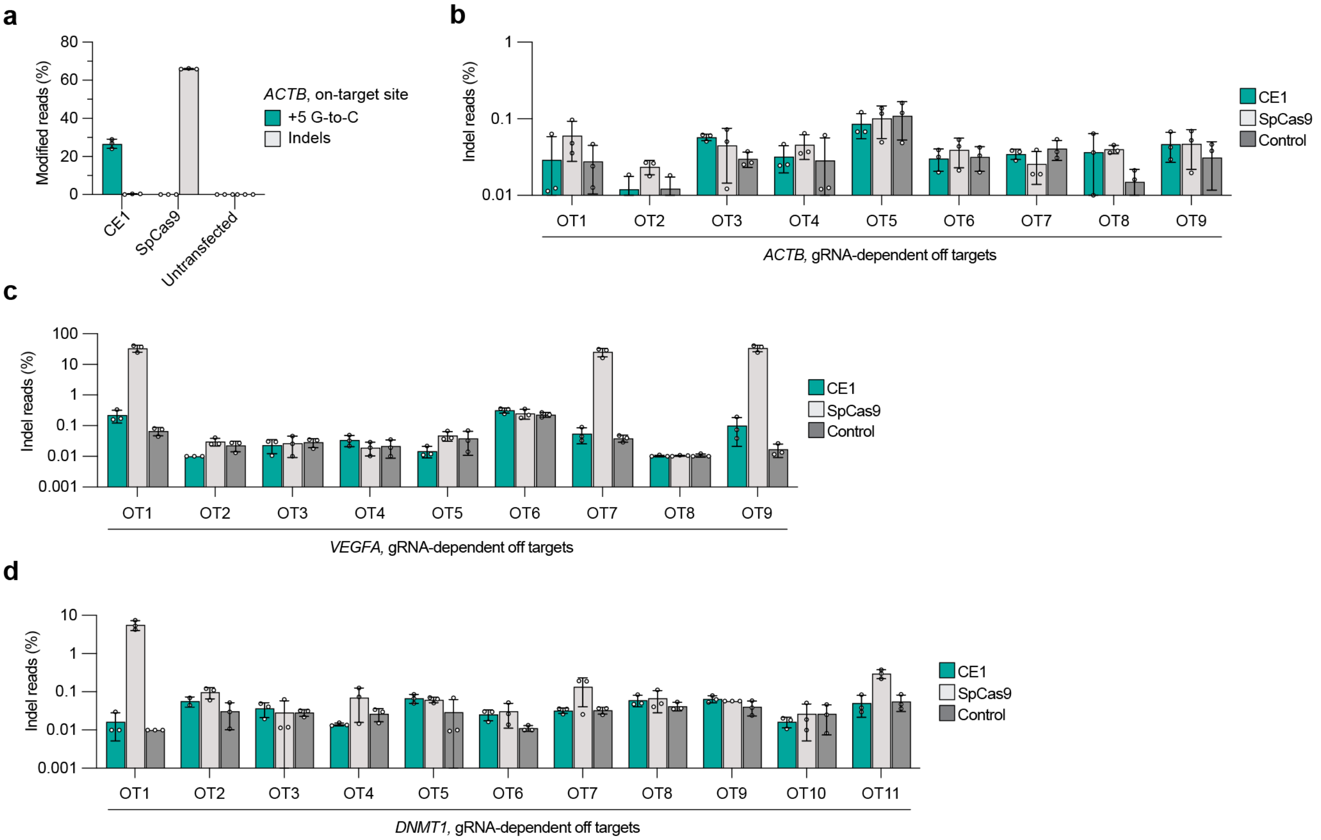


**Supplementary Figure 13. Cas9-dependent off-target characterization. a**, Percentage of sequencing reads harboring precise edits or indels at the *ACTB* on-target site, with CE1.n2b(+5) or SpCas9 nuclease. **b-d**, Percentage of sequencing reads harboring indels when using CE1 or SpCas9 nuclease at candidate off-target sites for gRNAs targeting *ACTB* (CE1.n2b(+5) for +5 G-to-C edit; **panel b**), *VEGFA* (CE1.n1 for a quadruple substitution edit; **panel c**), or *DNMT1* (CE1.n2(+49) for +3-5 delAGG edit; **panel d**). Putative off-target sites were nominated using Cas-OFFinder[13]. Data in all panels from HEK 293T cell experiments; mean, s.d., and individual datapoints shown for n = 3 independent biological replicates; Control data points were collected from genomic DNA extracted from untransfected cells.

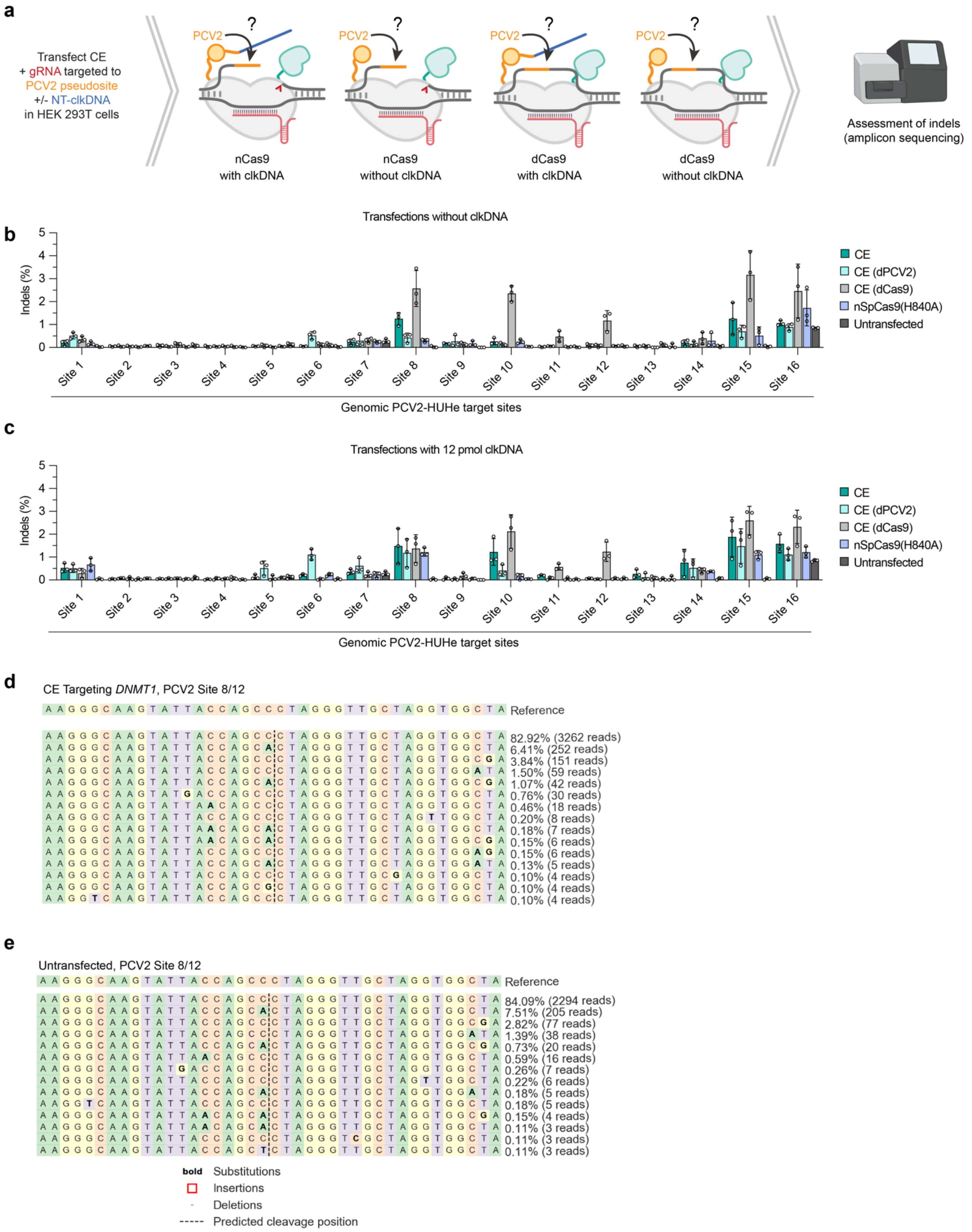


**Supplementary Figure 14. PCV2 HUHe off-target characterization.** **a**, Schematic of experimental conditions and workflow to characterize potential PCV2-mediated indels at genomic pseudosites bearing PCV2-HUHe binding motifs, when artificially inducing an R-loop via dCas9 or nCas9 binding. **b**,**c**, Percentage of sequencing reads harboring indels in the HUHe R-loop assay, quantified across 16 PCV2 pseudosites (**Sup. Note 2**), when transfected with the indicated constructs, a gRNA targeting the indicated HUHe pseudosite, and either with a clkDNA (**panel b**) or without clkDNA added (**panel c**). **d**,**e**, Representative allele frequency tables from CRISPResso2 (ref. [12]) outputs for site 8 or site 12 (the HUHe binding sites are in the same R-loop of separate spacers that are offset by 1 nt) when targeting a CE for productive click editing at *DNMT1* (**panel d**; see also **Fig. 4l**), or for an untransfected control (**panel e**; see also **Fig. 4l**). Data in **panels b** and **c** from HEK 293T cell experiments; mean, s.d., and individual datapoints shown for n = 3 independent biological replicates.

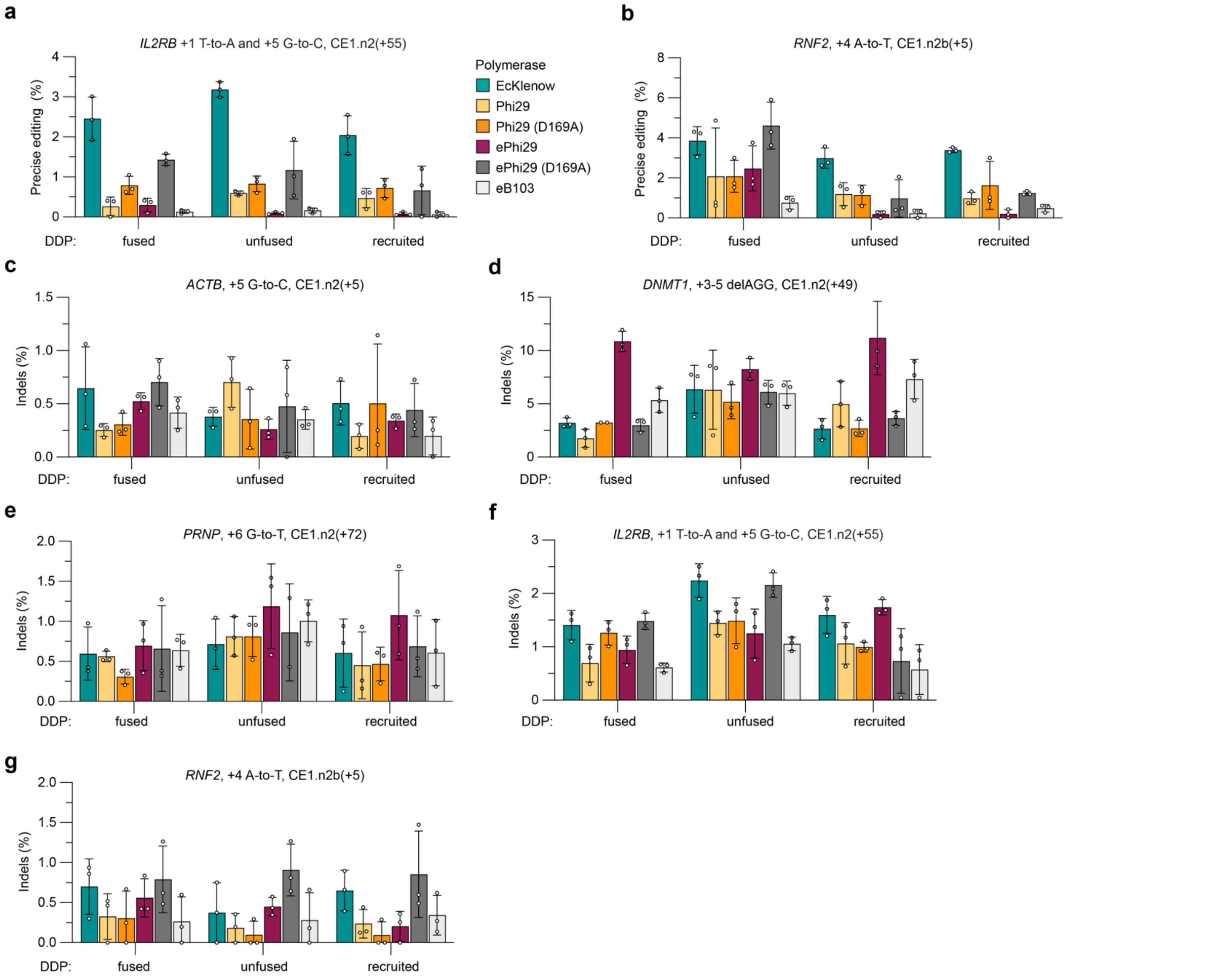


**Supplementary Figure 15. Characterization of CEs harboring different DDPs and architectures. a,b,** Percentage of sequencing reads with precise edits with gRNAs and clkDNAs targeting *IL2RB* (**panel a**) and *RNF2* (**panel b**) when using CE1.n2 constructs encoding different DNA-dependent polymerases and different construct architectures (fused, unfused or coiled-coil recruited). **c-g,** Percentage of reads with indels for gRNAs and clkDNAs targeting *ACTB*, *DNMT1*, *PRNP*, *IL2RB* and *RNF2* (**panels c-g**, respectively) when using CE1.n2 constructs encoding different DNA-dependent polymerases and different construct architectures; fused, covalent fusion of eHUH-nCas9-DDP; unfused, separately translated eHUH-nCas9 and DDP proteins; or recruited, where separately translated eHUH-nCas9 and DDP proteins have complementary N5/N6 coiled-coil peptides[14,15] fused to either protein). EcKlenow, Klenow fragment from *E.coli* DNA polymerase I (D355A, D357A); Phi29, DNA polymerase from bacteriophage ϕ29; Phi29 (D169A), 3'-5' exonuclease-deficient Phi29 DNA polymerase; ePhi29, engineered thermostable Phi29 DNA polymerase (M8R, V51A, M97T, G197D, E221K, Q497P, K512E, F526L); ePhi29 (D169), 3'-5' exonuclease-deficient ePhi29 (D169A, M8R, V51A, M97T, G197D, E221K, Q497P,

K512E, F526L); eB103, engineered thermostable Phi29 ortholog (H73R, A147K, R221Y, A318G, M339L, E359D, K372E, F383L, D384N, A503M, I511V, R544K, T550K. Data in **panels a-g** from HEK 293T cell experiments; mean, s.d., and individual datapoints shown for n = 3 independent biological replicates.
